# Structural and functional characterization of Rv0792c from *Mycobacterium tuberculosis*: identifying small molecule inhibitors against GntR protein

**DOI:** 10.1101/2021.09.17.460839

**Authors:** Neeraj Kumar Chauhan, Anjali Anand, Arun Sharma, Kanika Dhiman, Tannu Priya Gosain, Prashant Singh, Eshan Khan, Amit Kumar, Deepak Sharma, Ashish, Tarun Kumar Sharma, Ramandeep Singh

**Author notes:** **Corresponding author**: and. Translational Health Science and Technology Institute, Faridabad-Gurugram expressway, Faridabad-121001. and 91-129-2876370. Equal first author contributions (Neeraj Kumar Chauhan, Anjali Anand).

## Abstract

In order to adapt in host tissues, microbial pathogens regulate their gene expression through an array of transcription factors. Here, we have functionally characterized Rv0792c, a GntR homolog from *M. tuberculosis*. In comparison to the parental strain, ΔRv0792c mutant strain of *M. tuberculosis* was compromised for survival upon exposure to oxidative stress, cell wall agents and infection in guinea pigs. RNA-seq analysis revealed that Rv0792c regulates the expression of genes that are involved in stress adaptation and virulence of *M. tuberculosis*. Solution small angle X-ray scattering (SAXS) data steered model building confirmed that the C-terminal region plays a pivotal role in dimer formation. Systematic evolution of ligands by exponential enrichment resulted in identification of ssDNA aptamers that can be used as a tool to identify small molecule inhibitors targeting Rv0792c. Using SELEX and SAXS data based modelling, we identified residues essential for the DNA binding activity of Rv0792c and I-OMe-Tyrphostin as an inhibitor of Rv0792c aptamer binding activity. Taken together, we provide a detailed shape-function characterization of GntR family of transcription factors from *M. tuberculosis*. To the best of our knowledge, this is the first study that has resulted in the identification of small molecule inhibitors against GntR family of transcription factors from bacterial pathogens.

## INTRODUCTION

*Mycobacterium tuberculosis* (*M. tuberculosis*), the causative agent of Tuberculosis (TB) has coexisted with humans for thousands of years and is a leading cause of mortality among infectious diseases (1). Approximately, 2.0 million people are latently infected with *M. tuberculosis* due to the ability of the pathogen to persist in host tissues (2). The incidence rates of TB are on the rise due to HIV-TB coinfections, poor patient compliance, rise of drug-resistant strains and failure of BCG vaccine to impart protection against adult TB (3–6). Hence, there is a compelling need for identification of novel drug targets and regimens to tackle the problem imposed by primary and latent TB infections. *M. tuberculosis* is able to sense extracellular signals and reprogram its transcriptional profiles for stress adaptation and persistence in host tissues (7). This transcriptional reprograming is mediated by a complex network of regulatory proteins comprising of sigma factors, transcription factors, two components systems and serine-threonine protein kinases (8, 9). This highly coordinated regulation of gene expression in response to stress exposure is essential for *M. tuberculosis* to establish infection *in vivo*.

GntR family of transcription factors are highly abundant in various archaeal and bacterial genomes (10, 11). This protein family was named after the gluconate operon (gntRKPZ) repressor of *Bacillus subtilis* and is among the most widely distributed transcription factors in prokaryotes (12, 13). These proteins harbor a highly conserved amino-terminus DNA binding region and a carboxy terminus effector binding/oligomerization domain (10, 13). GntR proteins have an extended C-terminal which have not been visualized from any of the solved three-dimensional structure till date. Depending on the effector molecules, GntR family of transcription regulators have been categorized into various sub-families such as FadR, AraR, DevA, DasR, HutC, MocR, PlmA and YtrA (10, 13). Among these, FadR is the most well characterized GntR sub-family of transcription regulators. The binding of acyl-CoA to the effector domain of the FadR subfamily induces a conformational change and DNA binding is mediated by the helix-turn-helix motif (14). In addition to regulation of gluconate metabolism, GntR family of transcription factors have been shown to regulate microbial processes such as carbon metabolism, motility, antibiotic production, biofilm formation and pathogenesis (15–18). The genome of *M. tuberculosis* encodes for 8 homologs of GntR family of transcription regulators. Among these, Rv0043c, Rv0165c, Rv0494, Rv0586 and Rv3060c belongs to FadR family of GntR regulators. Previously, it has been shown that Rv0494 binds to fatty acyl CoA and negatively regulates the transcription of *kas* operon which is involved in mycolic acid biosynthesis (19). PipR (Rv0494 homolog) in *M. smegmatis* regulates the expression of genes involved in piperidine and pyrrolidine utilization of *M. smegmatis* (20). Rv0165c negatively regulates *mce1* operon and is necessary for *M. tuberculosis* persistence (21). Rv0586, negatively regulates the expression of *mce2* operon and endonuclease IV (22). In another study Zeng et al., demonstrated that the expression of genes required for vancomycin susceptibility are regulated by Rv1152 (23).

Here, we have functionally characterized Rv0792c from *M. tuberculosis* that belongs to HutC subfamily of GntR transcription regulators. We report that Rv0792c is an autoregulatory transcription factor and is required for *M. tuberculosis* survival in oxidative stress and to establish infection in host tissues. To the best of our knowledge, this is the first report where SELEX and SAXS approaches have been combined to determine: (i) the structural state of Rv0792c, (ii) aptamer binding pocket of Rv0792c and (iii) identification of I-OMe-Tyrphostin as a small molecule inhibitor of Rv0792c with a log IC_50_ value of ∼ 2.0µM. The results reported here are expected to pave ways to interfere or regulate the functioning of Rv0792c from *M. tuberculosis* to rationally identify and validate novel small molecule inhibitors.

## Materials and Methods

### Bacterial strains, media and growth conditions

The strains, plasmids and primers used in the study are shown in Table S1. Various *E. coli* strains were cultured in Luria-Bertani broth medium. *M. tuberculosis* H_37_Rv was used as a parental strain in this study. Unless mentioned, culturing of *M. tuberculosis* strains was performed in Middlebrook 7H9 and Middlebrook 7H11 medium as previously described (24). When required, the antibiotics were added at the following concentration; ampicillin, 100 μg/ml for *E. coli*, kanamycin, 25 μg/ml for both *E. coli* and mycobacteria, hygromycin, 150 μg/ml for *E. coli* and 50 μg/ml for mycobacteria and chloramphenicol, 34 μg/ml for *E. coli*. Biofilm experiments were performed in slightly modified Sauton’s medium: Ferric ammonium citrate 50 mg/L; MgSO_4_.7H2O 0.05 g/L; ZnSO_4_ 0.01 mg/L; K2HPO_4_ 1.0 g/L, CaCl_2_ 0.05 g/L; asparagine 0.5 g/L; Na_2_HPO_4_ 2.5 g/L and Tyloxapol 0.05%. MIC_99_ determination assays were performed using microdilution method in the presence of different drugs as described previously (25).

### Protein expression and purification

DNA fragments coding for either wild type Rv0792c or mutant proteins (Rv0792c^R49A^ or Rv0792c^G80D^ or Rv0792c^R41A^ or Rv0792c^P40A^) were cloned into IPTG inducible prokaryotic expression vector, pET28b. For protein expression and purification, various constructs were transformed into *Escherichia coli* BL21-CodonPlus strain. The expression of recombinant proteins was induced by the addition of 1.0 mM IPTG at 18 °C at 200 rpm for 12-16 hrs. The induced cultures were harvested by centrifugation at 6000*g*, pellets were resuspended and clarified lysates were prepared by sonication in 1x PBS. The recombinant protein from the clarified lysates was purified using Ni^2+^-NTA chromatography. The purified fractions were analysed by SDS-PAGE, pooled, dialyzed, concentrated and stored at −80°C in the buffer (10 mM Tris - pH 7.4, 100 mM NaCl and 5% glycerol) till further use.

### Sedimentation velocity experiments

Analytical ultracentrifuge experiments were performed in Beckman optima XL-I analytical ultracentrifuge equipped with an absorbance-based detection system. The two-sector charcoal centrepiece of 1.2 cm path length built with sapphire windows was used for the study. The cell was filled with 390 µL of protein sample at a concentration of 0.38 mg/mL, 0.76 mg/mL or 1.52 mg/mL in Buffer A (25 mM HEPES - pH 7.2, 400 mM NaCl and 10% Glycerol). The reference cell was filled with 400 µL of buffer A. Radial scans were collected at 40,000 rpm for 12-14 hours with an interval of 3 mins at 280nm. The data analysis was performed with the ‘SedFit analysis’ program using continuous distribution(s) model based on the Lamm equation. Parameters such as buffer density (ρ=1.01660), buffer viscosity (η=0.01057 poise) and partial specific volume (ʋ=0.73677) of protein were measured by the Sednterp program. Standard sedimentation coefficients of samples were represented as S_20,w_ referring sedimentation at 20°C.

### Generation of Rv0792c mutant and complemented strain of *M. tuberculosis*

In order to investigate th e role of Rv0792c in *M. tuberculosis* physiology and pathogenesis, the mutant strain was constructed using temperature sensitive mycobacteriophages as per standard protocols (26). The replacement of the Rv0792c coding region with the hygromycin resistance gene in the mutant strain was confirmed by PCR and qPCR using locus specific primers. For the generation of complemented strain, Rv0792c was amplified along with its native promoter and cloned into an integrative vector, pMV306K. The electrocompetent cells of Rv0792c mutant strain was electroporated with the recombinant plasmid pMV306K-Rv0792c and the transformants were selected on Middlebrook 7H11 medium supplemented with kanamycin and hygromycin.

### Stress Experiments

In order to evaluate the role of Rv0792c in stress adaptation, early-log phase cultures (OD_600nm_ ∼ 0.2) of various strains were subjected to different stress conditions as previously described (24, 27). For drug tolerance experiments, mid-log phase cultures (OD_600nm_ ∼ 1.0) were exposed to different drugs such as isoniazid, rifampicin and levofloxacin at 10x MIC_99_ concentration. For bacterial load enumeration, 10-fold serial dilutions were prepared and 100 μl was plated on Middlebrook 7H11 medium at 37°C for 3-4 weeks.

### Animal Experiments

For animal experiments, pathogen-free female guinea pigs (Hartley strain weighing, 250–300 g) were purchased from Lala Lajpat Rai University of Veterinary and Animal Sciences, Hisar, India. The infection experiments were supervised as per CPCSEA guidelines and performed at the Infectious Disease Research Facility, Translational Health Science and Technology Institute, New Delhi, India. The animal experiments were approved by the Institutional animal ethics committee of Translational Health Science and Technology Institute. For aerosol infection, the strains were grown till mid-log phase, washed twice with 1x PBS and single cell suspensions were prepared. The guinea pigs were infected using a Glas-Col aerosol chamber with single cell suspensions of various strains that resulted in implantation of 50-100 bacilli at day 1 post-infection. The extent of disease progression in guinea pigs was determined by both CFU and histopathology analysis as described previously (24).

### RNA-seq analysis

For RNA-seq analysis, total RNA was isolated from mid-log phase (OD_600nm_ ∼ 0.8) cultures of parental and Rv0792c mutant strain using Trizol method as described previously (24). The purified RNA was shipped to Aggrigenome Labs Pvt Ltd (India) for sequencing. The preparation of library, RNA-sequencing and data analysis was performed as previously described (28). The transcripts showing differential expression of more than 2.0 fold with a *P-value* <0.05 were considered to be significant. Quantitative PCR was performed to validate the identified DEGs in the Rv0792c mutant strain. For qPCR studies, cDNA was synthesized using 200 ng of DNase I treated mRNA and Superscript III reverse transcriptase as per manufacturer’s protocols. The synthesized cDNA was diluted 1:5 and qPCR was performed using gene specific primers and SYBR green mix. The expression of genes of interest was normalized to the transcript levels of *sigA* (a housekeeping gene) and was quantified as previously described (24).

### Aptamer selection against Rv0792c

Aptamer selection against Rv0792c was performed using SELEX method as previously described (29). Briefly, to identify unique aptamer sequences specific for Rv0792c, aptamer library (2000 picomoles) in selection buffer (SB; 10 mM Tris, pH-7.5, 10 mM MgCl_2_, 50 mM KCl, 25 mM NaCl) was allowed to bind nitrocellulose membrane (NCM) to remove the NCM binding species. The unbound ssDNA was removed from the NCM and incubated with (His)_6_-Rv0792c immobilized membrane. After one hour of incubation, the membrane was washed with SB supplemented with 0.05% Tween-20. The protein bound aptamers were eluted by heating at 92°C and further enriched through PCR. The stringency of selection in successive rounds was enhanced by (i) increasing the number of washes (2-5 times), (ii) the strength of buffer (0.05-1% Tween-20), (iii) decreasing incubation time in the selection step and (iv) increasing the incubation time for negative selection steps. After 6 rounds of selection, the highest affinity pool was cloned in pTZ57R/T vector. The plasmid DNA was isolated from randomly picked transformants and confirmed by sequencing. Subsequently, the sequence homology among the sequenced aptamers was performed using CLUSTALW and Bio Edit sequence alignment editor.

### Aptamer Linked Immobilized Sorbent Assay (ALISA) and determination of dissociation constant (K_d_)

ALISA was performed as previously described (29). Briefly, a Nunc MaxiSorp™ 96 well-plate was coated with 500 ng of purified wild type or mutant proteins in 100 µL (0.1 mol/L) sodium bicarbonate buffer (pH-9.6) at 37°C for 2 hrs. The plate was incubated with blocking buffer (5% BSA and 0.25% Tween-20) at room temperature. After blocking, the wells were washed once with SB. This was followed by the addition of 100 pmol of 5′ biotinylated aptamer in SB and plates were further incubated at room temperature. Following incubation for 1 hr, the wells were washed twice with SB supplemented with 1% Tween-20 and SB only. Following this, 1:3000 streptavidin-horse radish peroxidase antibody (strep-HRP) was added for 1 hr at room temperature. The unbound antibody was removed by washing and 100 µL of 3, 3′, 5, 5′-tetramethylbenzidine (TMB) was added. The reaction was stopped after 5-10 mins of incubation by the addition of 100 µL of 5% H_2_SO_4_. The quantification of the protein-bound aptamer-strep complex was determined by measuring absorbance at 450 nm. For K_d_ determination, 500 ng protein was coated per well. After blocking, aptamer was added in the range of 2-500 nM and ALISA was performed as described above. Next, the absorbance reading at 450nm was plotted as a function of aptamer concentration and K_d_ was determined using non-linear regression model in Graph-pad Prism.

### Electromobility gel shift assay (EMSA) for aptamer and promoter binding with Rv0792c

For EMSA, 100 pmol of selected aptamer candidates were incubated with 8 µM and 16 µM of Rv0792c protein in binding buffer (10 mM Tris, pH-7.8, 10 mM MgCl_2_, 50 mM KCl, 25 mM NaCl, 0.5 mg salmon sperm DNA, 0.75 mg BSA and 2.5% glycerol) for 20 mins at 4°C. The reactions were resolved on 2% agarose gel and the protein-aptamer complexes were detected by staining using SYBR safe. For promoter binding assays, the amplified fragments were gel extracted, end-labeled with [γ-^32^P] ATP (1000 Ci nmol^−1^) using T4 polynucleotide kinase and purified using Sephadex G-50 spin columns (30). Promoter binding assays were performed using radiolabeled promoter fragment and 1 μM of (His)_6_-Rv0792c in binding buffer (10 mM Tris, pH-7.4, 50 mM NaCl, 10 mM MgCl_2,_ 0.2 mg/ml BSA, 10% glycerol, 1 mM dithiothreitol (DTT), and 200 ng of sheared herring sperm DNA). Incubation was performed on ice for 20 mins and subsequently, the reactions were resolved on 6% non-denaturing polyacrylamide gels at 4°C, dried and bands were visualized using a phosphorimager.

### Small angle X-ray scattering (SAXS) AND Model Building

The SAXS studies were performed at an in-house SAXspace instrument using a line collimation source (X-rays wavelength 0.15414 nm) (Anton Paar, Graz, Austria). The samples were studied with sample to detector distance of 317.06 mm. About 50 μL of unliganded proteins and aptamers, their molar mixtures and matched buffer were transferred in a 1 mm thermostated quartz capillary, and scattered X-rays were monitored on a 1D Mythen detector. An average of three frames of one hour each was obtained for further data analysis. Table S2 mentions the programs used for different steps of data collection and processing. SAXS data analyses were performed using the ATSAS suite of programs v 2.8.4. Using shape constraints in the SAXS data profile, the shape of the predominant scattering particles was restored. Residue details models of unliganded proteins/aptamers and their docked structures were compared and superimposed using automated alignment of inertial axes using CRYSOL and SUPCOMB programs in the suite, respectively.

The models of unliganded (His)_6_-Rv0792c and ssDNA aptamers with residue/base level details were generated using the primary structures of protein and aptamers. SWISS-MODEL server was used to search for structural templates of Rv0792c (https://swissmodel.expasy.org) (31). The results provided the putative template for Rv0792c from residues 53-286 of lin2111 from *Listeria innocua* [PDB ID: 3EDP]. The amino-terminus 1-52 residues and carboxy terminus segment from 287-303 residues were generated using molecular dynamics of the segments. MD stimulation studies were performed using Tinker molecular modeling package v 4.2 along with OPLSUA forcefield. Advanced Newton Raphson method was employed to compute structures of segments at 298K in implicit water (ε = 80). Simulations were run for 10 ns with restart coordinates written at every 1 ps. The predominant low energy structures were filtered out as described previously (32, 33). SAXS data supported a dimeric state of unliganded Rv0792c, thus using the dimer as a central scaffold, two copies of predominant low energy conformations of N- and C-terminal segments were aligned in space using the SASREF program, as reported previously (34). The composite structure of Rv0792c was generated and energy minimized by performing template-based modeling using the SWISS-MODEL server. ELNEMO server was used to compute low frequency collective vibrations accessible to the protein structure (35). ssDNA aptamers were modeled using their sequences and the ICM 3.8 program. SAXS data supported their masses to be close to their monomers and thus monomeric forms of all three aptamers were considered for modeling studies. Repeated runs of global minimization and local optimization were performed till the conformations did not change more than 0.01 RMSD across all atoms. By comparing theoretical SAXS profiles of the ten lowest energy conformations of aptamers with experimental SAXS data on the aptamers, the best conformation of aptamer agreeing with experimental data was identified. Further, the ELNEMO server was performed to compute the most collective low energy vibration mode to compare with SAXS data-based information and shape. These structures of ssDNA aptamers were docked on the structure of dimeric Rv0792c protein, and their pose on the protein was identified using SAXS data of the complexes as reference. The graphs pertaining to SAXS data analysis were prepared using OriginLab v5 software. The images of molecular models were prepared using open-source Pymol v 1.1 and UCSF Chimera softwares v 1.14.

### *In silico* screening of drug-like molecules

Molecular docking studies were performed using ICM Chemist Pro software v3.8. Using the interaction distance mapping option, all residues within 3 Å of the interacting surface of Rv0792c to aptamers, from the models of protein: aptamer complexes were selected. From both chains of Rv0792c, stretches of residues 125-137, 224-231, 253-257, 279-289, and 300-303 were selected to form the aptamer binding site. The library of approved drugs from drugbank.ca was used for docking studies, and the docking was done in an automated manner. Full degrees of freedom and rotations were given to the ligand during evaluating docking poses on the identified receptor surface. The docking was carried out individually with each ligand, and its various poses with respective to the receptor pocket’s charge and shape profile was calculated. The scores obtained of the docked pose were then arranged from low to high, and the top ten lowest scoring ligands were further selected for optimizing receptor residues around the low score pose of ligand to obtain new score. The shortlisted ligands were further arranged according to the 4D docking score.

Next, we performed competitive ALISA to determine the ability of top two hits to compete with the binding of aptamer to Rv0792c. The coating of the wild-type protein and blocking of non-specific sites was performed as described above. Subsequently, the binding of aptamer was determined in the presence or absence of the top-two small molecule inhibitors. For IC_50_ determination assays, inhibition assays were performed in the presence of 2.0-fold serial dilutions of small molecules. IC_50_ values were calculated as the drug concentration that showed 50% inhibition for aptamer binding with Rv0792c.

### Data availability

The raw data files for RNA-seq experiments has been deposited at NCBI: PRNJA727912. The data pertaining to unliganded protein, aptamer and protein-aptamer complexes is available at https://www.sasbdb.org/project/1396/kv4wukfdsj.

### Statistical analysis

GraphPad Prism 8 software (version 8.4.3, GraphPad Software Inc., CA, USA) was used for statistical analysis and graphs generation. Significant differences between indicated groups were calculated using the ‘t-test’ function and were considered significant at a *P*-value of <0.05.

## RESULTS

### Rv0792c from *M. tuberculosis* belongs to HutC sub-family of GntR transcription factors

GntR family of transcription factors are highly conserved in the bacterial kingdom and *M. tuberculosis* genome encodes for eight GntR homologs (Rv0043c, Rv0165c, Rv0494, Rv0586, Rv0792c, Rv1152, Rv3060c and Rv3575c). Multiple sequence alignment revealed that GntR homologs from *M. tuberculosis* shared almost identical residues in DNA binding amino terminus region. As expected, not much sequence identity was seen in the effector binding region of the GntR family of transcription regulators. Phylogenetic analysis revealed the formation of two preponderant groups. Group-1 is the largest and ∼91% of proteins are clustered together in Group-1, possibly due to high similarity in amino acid sequences (Fig. 1A). As shown in Fig. 1A, Rv0792c clustered with different proteins of the GntR family such as HutC from *P. putida*, DasR from *S. coelicolor*, NagR from *B. subtilis*, PhnR from *S. enterica*, MngR from *E. coli* (10, 36–38). Multiple sequence alignment analysis between Rv0792c and other HutC homologs revealed that these proteins share an identity of ∼ 30% among themselves (Fig. S1). FadR homologs from *M. tuberculosis* (Rv0494, Rv0586 and Rv3060c) and YtrA homolog (Rv1152) also grouped with their respective analogs from other bacterial species (Fig. 1A). The Group-2 cluster consisted of Rv3575c and GntR homolog from *Klebsiella Pneumoniae* that shared similarity 24% among themselves (Fig. 1A). In the present study, we have performed experiments to biochemically, functionally and structurally characterize Rv0792c from *M. tuberculosis*.

**Figure 1:**
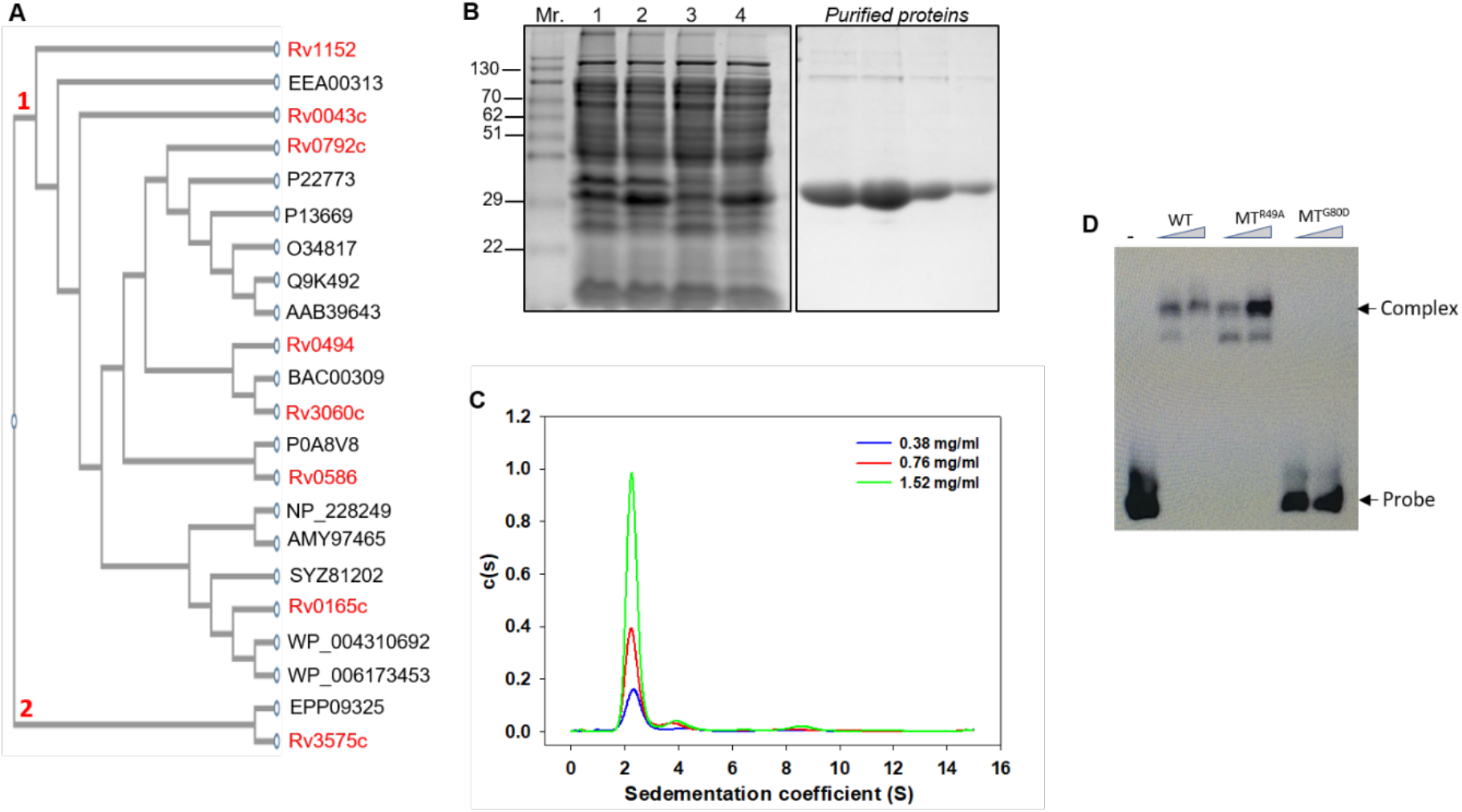
**(A) Rooted phylogenetic tree for GntR family of transcription factor in various prokaryotes.** The bacterial species included in the phylogenetic analysis are: *M. tuberculosis* (Rv0043c, Rv0165c, Rv0494, Rv0586, Rv1152, Rv0792c and Rv3060c), Burkholderia sp (EEA00313), *Pseudomonas Putida* (P22773), *Escherichia coli* (P0A8V8, P13669), *Bacillus subtilis* (O34817), *Streptomyces coelicolor* (Q9K492), *Salmonella enterica* (AAB39463)*, Corynebacterium glutamicum* (BAC00309)*, Thermotoga maritima* (NP_228249), *Streptococcus pyogenes* (AMY97465), *Vibrio cholerae* (SYZ81202), *Pseudomonas syringae* (WP_044310692), *Brucella sp* (WP_006173453) and *Klebsiella pneumoniae* (EPP09325). **(B) Rv0792c expression and purification.** SDS-PAGE analysis of Rv0792c expression in *E. coli*. Mr, molecular size markers (molecular weight of Rv0792c is about 29.9 kDa); Lane 1, Uninduced total fraction; Lane 2, Induced total fraction; Lane 3, Uninduced soluble fraction; Lane 4, Induced soluble fraction. Protein fractions purified by Ni-NTA affinity chromatography. **(C) Sedimentation velocity analytical ultracentrifugation experiments.** Radial absorption curves for different concentrations of Rv0792c were collected at 280nm. Scans were collected every 3 mins. Sedimentation coefficient continuous distribution c(s) plots of the protein at indicated concentrations. c(s) plots show that Rv0792c exists majorly as dimer in its native state. **(D) Rv0792c binds to its own promoter.** EMSAs of (His)_6_-Rv0792c protein with the promoter region of Rv0792c. The labelled probe was incubated with either wild type or various mutant proteins as outlined in Materials and Methods.

### Rv0792c is a dimeric protein and bind its own promoter

For biochemical characterization, Rv0792c was cloned in pET28b and recombinant protein was purified with amino-terminus histidine tag (Fig. 1B). The purity of various fractions was confirmed by SDS-PAGE analysis. The purified fractions were dialyzed, concentrated and subjected to sedimentation velocity ultracentrifugation experiments at varying protein concentrations, 0.38 mg/mL, 0.76 mg/mL and 1.52 mg/mL (Fig. 1C). The continuous distribution (c(s)) analysis of absorbance scans at different protein concentrations revealed that Rv0792c predominantly sediments at s_20,w_ of ∼2.3S, consistent with a molecular weight of ∼58kDa, thereby suggesting the protein is primarily dimeric in solution (Fig. 1C and Table 1). Additionally, very small fractions of higher order oligomeric species sedimenting at s_20,w_ of ∼4.3S (8-11%) and ∼8.5S (4-5%) corresponding to 130 kDa and 396 kDa, respectively, were also observed (Fig. 1C). The increase in fraction of species sedimenting at ∼4.3S and ∼8.5S with increased protein concentration suggest formation of higher order oligomers at relatively higher concentrations of the protein.

**Table 1.**
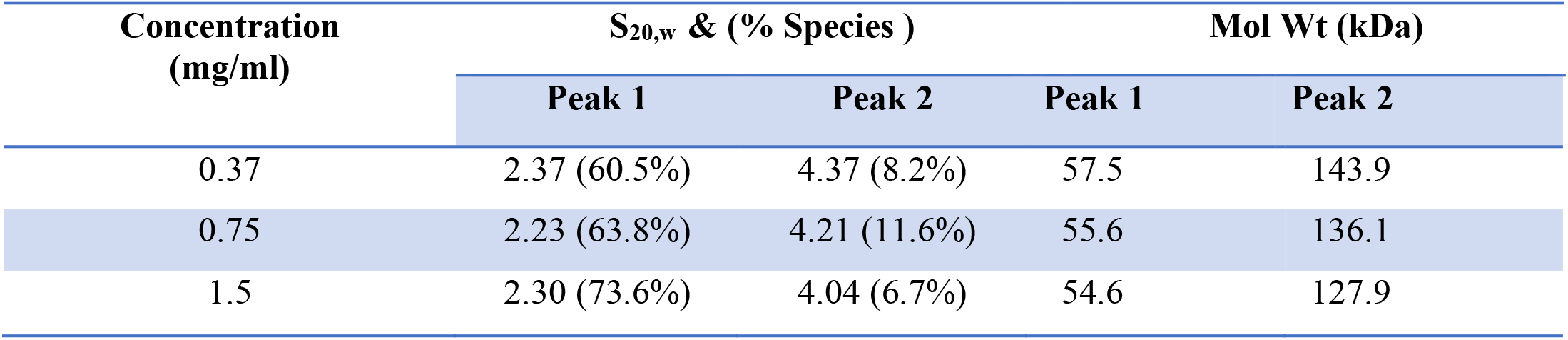
Sedimentation coefficient values of Rv0792c obtained from analytical ultracentrifugation studies.

| Concentration<br>(mg/ml) | S<sub>20,w</sub> & (% Species ) |  | Mol Wt (kDa) |  |
| --- | --- | --- | --- | --- |
|  | Peak 1 | Peak 2 | Peak 1 | Peak 2 |
| 0.37 | 2.37 (60.5%) | 4.37 (8.2%) | 57.5 | 143.9 |
| 0.75 | 2.23 (63.8%) | 4.21 (11.6%) | 55.6 | 136.1 |
| 1.5 | 2.30 (73.6%) | 4.04 (6.7%) | 54.6 | 127.9 |

It has been previously reported that the GntR family of transcription factors bind to their own promoters and autoregulate their own expression (39, 40). Therefore, we next performed EMSA assays to study the binding of purified Rv0792c with its native promoter. As shown in Fig. 1D, the purified protein was able to bind to the radiolabeled promoter in a dose-dependent manner. Clear retardation was seen in the mobility of labeled DNA in the presence of purified protein. These observations suggest that similar to other GntR homologs, Rv0792c binds to its own promoter and likely autoregulates its expression (39, 40). DNA binding domains of GntR protein at the amino-terminus are highly conserved (14, 41). Multiple sequence alignment analysis revealed that residues important for DNA binding, Arg49 and Gly80 of Rv0792c that corresponds to Arg35 and Gly66 of FadR proteins were conserved. Next, we performed EMSA assays using labeled Rv0792c promoter and purified wild type, Rv0792c^R49A^ and Rv0792c^G80D^ mutant proteins. We observed that Rv0792c^R49A^ mutant protein binds to the Rv0792c promoter (Fig. 1D). However, the mutation of glycine 80 to aspartic acid completely abrogated the ability of Rv0792c to bind to its native promoter (Fig. 1D). Based on these findings, we conclude that Rv0792c binds to its own promoter and glycine residue at position 80 is essential for its DNA binding ability.

### Rv0792c is essential for the adaptation of *M. tuberculosis* upon exposure to oxidative and cell wall damaging agent

TB infection is an outcome of *M. tuberculosis* adaptation to unfavorable environmental conditions encountered in host tissues such as low oxygen, nutrient limitation, reactive nitrogen intermediates, oxidative and acidic stress (42). Several metabolic pathways including transcriptional regulators are essential for *M. tuberculosis* pathogenesis. The exact role of GntR homologs in *M. tuberculosis* pathogenesis has not been deciphered extensively. Here, we determined the role of Rv0792c in physiology, stress adaptation and virulence of *M. tuberculosis*. Using temperature sensitive mycobacteriophages, we generated a Δ0792c mutant strain of *M. tuberculosis* (Fig. S2A). The construction of the mutant strain was confirmed by PCR and qPCR using gene specific primers. As shown in Fig. S2B, locus specific primers resulted in amplification of ∼ 1kb and 2.1 kb bands in the case of wild type and mutant strain, respectively. The restoration of Rv0792c expression in the complemented strain was confirmed by qPCR (Fig. S2C). As shown in Fig. S2D and S2E, no changes were observed in growth patterns and colony morphology of parental and Δ0792c mutant strain of *M. tuberculosis*.

We next compared the survival of various strains in different stress conditions *in vitro*. As shown in Fig. 2A, a growth defect of 5.0- and 11.0-folds was seen in the survival of mutant strain in comparison to the parental strain after exposure to oxidative stress for 24 hrs and 72 hrs, respectively (Fig. 2A, \**P<0.05*). This growth defect associated with the mutant strain was restored in the complemented strain (Fig. 2A). The mutant strain also exhibited a ∼ 5.0-fold growth defect after exposure to cell wall degrading agent, lysozyme in comparison to the wild type strain (Fig. 2B, *\*\*P<0.01*). We observed that both wild type and mutant strains were susceptible to comparable levels after exposure to other stress conditions tested in this study (Fig. 2C, 2D, 2E and 2F). Since GntR’s have been shown to be involved in the biofilm formation of bacterial pathogens, we also determined the role of Rv0792c in *M. tuberculosis* biofilm formation *in vitro* (43–46). We observed that the parental and Δ0792c mutant strain of *M. tuberculosis* were comparable in their ability to form biofilms *in vitro* (data not shown). In order to understand the role of Rv0792c in drug tolerance, we next determined the susceptibility of various strains to drugs with a different mechanism of action. Fig.S2F unequivocally demonstrates that deletion of Rv0792c did not alter the susceptibility of *M. tuberculosis* after 14 days of exposure to aforesaid drugs. In concordance, both strains displayed comparable MIC_99_ values of 0.39 μM, <0.05 μM, 0.78 μM and 3.125 μM against isoniazid, rifampicin, levofloxacin and ethambutol, respectively. Taken together, we demonstrate that Rv0792c is important for the adaptation of *M. tuberculosis* upon exposure to oxidative stress *in vitro*.

**Figure 2:**
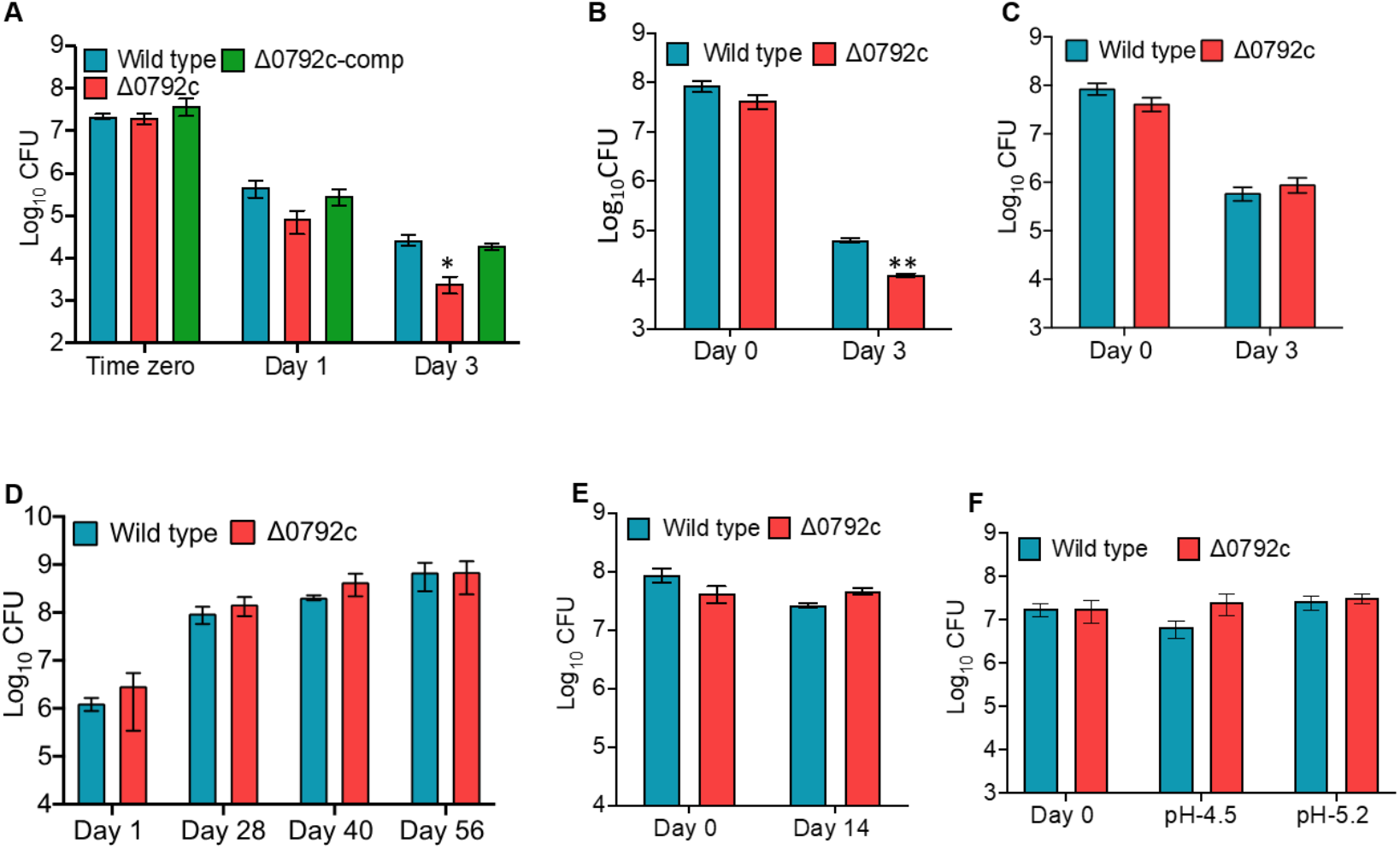
The effect of deletion of Rv0792c on the survival of *M. tuberculosis* in different stress conditions *in vitro.* For *in vitro* stress experiments, early log phase cultures of various strains were exposed to either oxidative stress (5 mM H_2_O_2_, A) or lysozyme (2.5 mg/ml, B) or nitrosative (5mM NaNO_2_, C) or low oxygen (D) or nutritional stress (1x TBST, E) or acidic pH (pH-5.2, F). The data shown in these panels is mean ± S.E. of log_10_ CFU obtained from two or three independent experiments. The statistically significant differences were observed for the indicated groups (paired (two-tailed) *t*-test, \**P*<0.05, *\*\*P<0.01*.

### Deletion of Rv0792c impairs the ability of *M. tuberculosis* to cause infection in guinea pigs

Next, we determined the ability of Rv0792c to contribute to *M. tuberculosis* pathogenesis using the guinea pig model of infection. For animal experiments, guinea pigs were infected with various strains and aerosol infection resulted in implantation of ∼50-100 bacilli in lungs at day 1 post-infection. We observed discrete multiple lesions in lung tissues of animals infected with parental and complemented strain. The number of these lesions were significantly reduced in lung tissues from mutant strain infected guinea pigs at both 28 - and 56-days post-infection (Fig. 3A). We did not observe any differences between the weight of lung tissues of guinea pigs infected with various strains. As shown in Fig. 3B, at 4 weeks post-infection ∼ 230.0-fold significantly higher bacterial numbers were observed in lungs of wild type strain infected guinea pigs in comparison to the mutant strain infected guinea pigs (*\*\*P<0.01*). In agreement, splenic bacillary loads were higher in guinea pigs infected with wild type strain by ∼ 57.5-fold compared to the mutant strain infected animals (Fig. 3C, \**\*P<0.01*). Further, the lungs and splenic bacillary loads were reduced by ∼ 120.0- and ∼ 600.0-folds, respectively, in mutant strain infected guinea pigs, in comparison to wild type infected guinea pigs at 56-days post-infection (Fig. 3D, 3E, \**\*\*P<0.001, **P<0.01*). We also observed that complementation with Rv0792c partially restored the growth defect associated with the mutant strain at both 4- and 8-weeks post-infection (Fig. 3B, 3C, 3D and 3E). Concordantly, minimal tissue involvement was observed in hematoxylin and eosin stained sections from guinea pigs infected with the mutant strain at 8 weeks post-infection (Fig. 3F). The granuloma formation was seen in sections from animals infected with either wild-type or complemented strains (Fig. 3F). Taken together, we show that Rv0792c is not essential for growth *in vitro* but is indispensable for *M. tuberculosis* to establish infection in host tissues.

**Figure 3:**
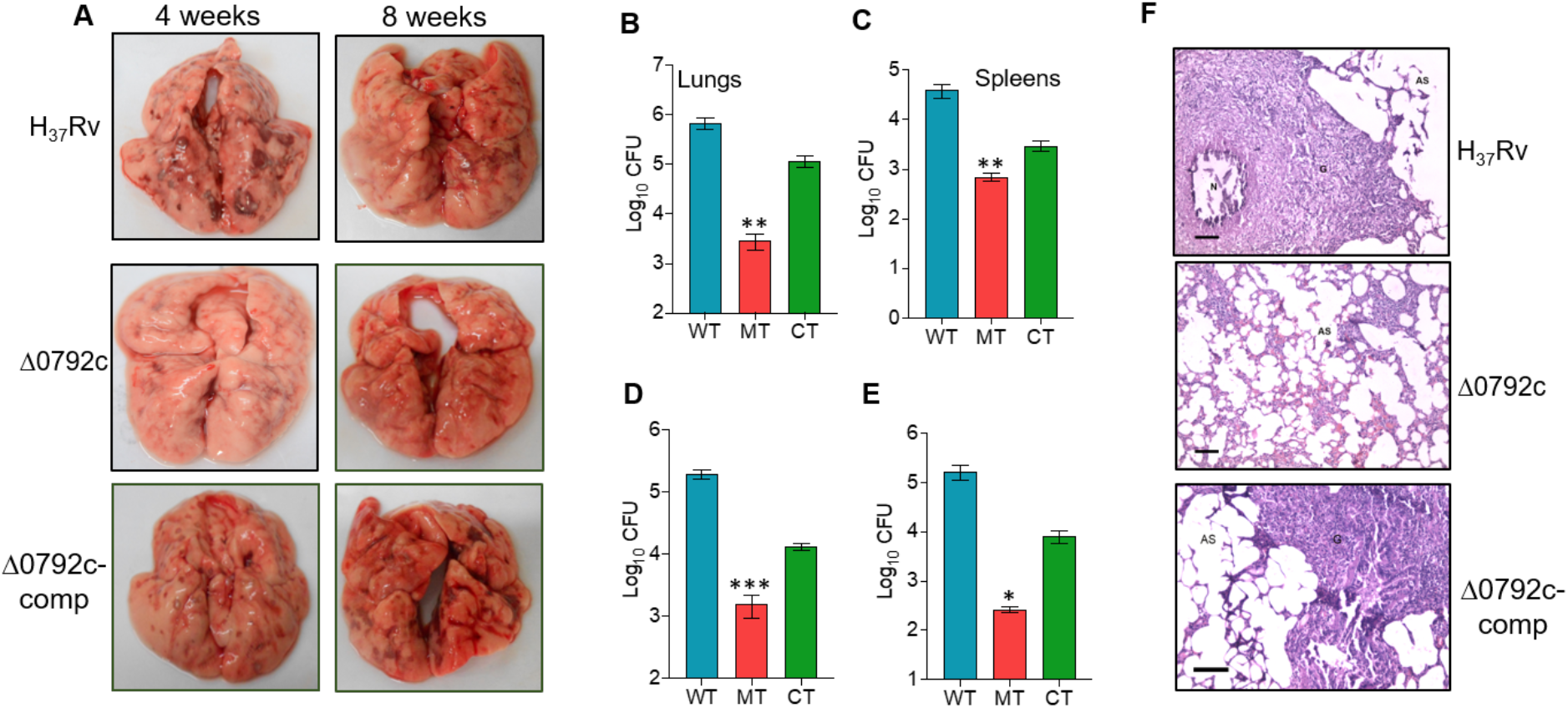
Rv0792c is essential for *M. tuberculosis* to establish infection in the host. **(A)** Gross pathological evaluation of tissue damage of lung tissues from guinea pigs infected with either wild type or Rv0792c mutant or Rv0792c complemented strains at 4 weeks or 8 weeks post-infection. **(B-E)** The lung and splenic bacillary loads were determined in aerosol infected guinea pigs at both 4 weeks (B, C) and 8 weeks (D, E) post-infection. The data represented in this panel is mean ± S.E. of log_10_ CFU obtained from 6 or 7 animals per time points for each strain. The statistically significant differences were observed for the indicated groups (paired (two-tailed) *t*-test, *\*P<0.05*, \*\**P<0.01*, *\*\*\*P<0.001*. **(F)** The tissue sections from 8 weeks infected animals were stained with haematoxylin and eosin to determine the extent of tissue damage. Scale bar, 100 microns, 100x magnification.

### Effect of deletion of Rv0792c on the transcriptional profile of *M. tuberculosis*

The observed growth defect of the Rv0792c mutant strain in guinea pigs suggests that it might regulate the expression of genes that are involved in the virulence or stress adaptation of *M. tuberculosis*. In order to define Rv0792c regulon, RNA-seq experiments were performed using total RNA isolated from mid-log phase cultures of wild type and the mutant strain as described in Materials and Methods. We observed that majority of the genes were expressed to similar levels in the mutant and wild type strain. Using a cut-off value of >2.0-fold change and *p-value <0.05*, transcriptome analysis revealed that a total of 197 genes were differentially expressed in mutant strain compared to the wild type strain (Fig. 4A). Among these, the levels of 108 and 89 transcripts were increased and decreased, respectively, in the mutant strain (Fig. 4A, Table S4). These differentially expressed genes were further characterized based on their annotations in Mycobrowser (https://mycobrowser.epfl.ch/). We noticed that most of the differentially expressed genes were either conserved hypothetical proteins or involved in processes such as cell wall synthesis or intermediary metabolism (Fig. 4B). The transcript levels of proteins such as Rv0383c, Rv1094 (*desA2*), Rv1285 (*cysD*), Rv1350 (*fabG2*), Rv2166c, Rv2846c, Rv2988c and Rv3139 that are essential for *M. tuberculosis* growth *in vitro* were downregulated in the mutant strain (Table S4). RNA-seq analysis revealed that the transcripts of genes upregulated in low oxygen conditions (such as Rv2624c, Rv2625c, Rv3126c, Rv0572c and Rv1734c) or nutrient limiting conditions (such as Rv1149, Rv1285, Rv1929c, Rv2169c, Rv2269c, Rv2660c and Rv2745c) were reduced in the mutant strain (Table S4, (47, 48). Among, the upregulated genes, the transcript levels of genes adjacent to Rv0792c and *mymA* operon were increased in the mutant strain (49, 50). A subset of these differentially expressed genes in RNA-seq experiments was also assessed by qPCR. As expected, the expression patterns obtained by qPCR were similar to those obtained from RNA-seq data (Fig. 4C). These observations indicate that Rv0792c regulates the expression of genes and this transcriptional reprogramming is required for *M. tuberculosis* to adapt and survive in host tissues.

**Figure 4:**
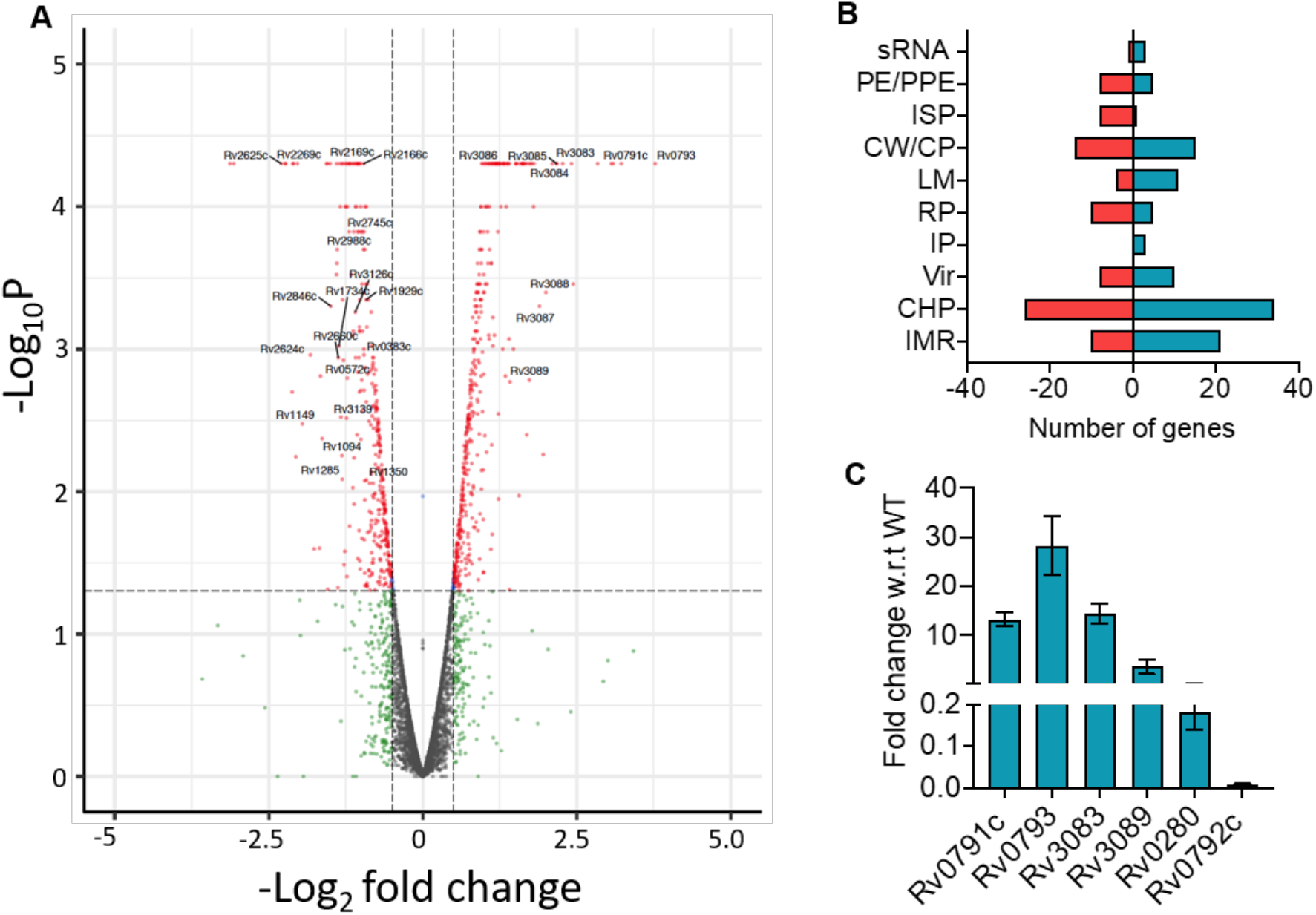
Differential expression of genes in Rv0792c mutant strain. **(A)** Volcano plot illustrating the gene expression comparison between wild type vs mutant strain, where individual genes are represented as single dots with the -log_10_ *P*-value on the y axis and the log_2_ fold change on the x axis. Selected subset of significant genes are shown with their gene names as the labels. **(B)** The functional annotation of differentially expressed genes between parental and Rv0792c strain is shown. These annotations were generated using TB data base (https://mycobrowser/epfl.ch). The upregulated and downregulated genes in the mutant strain has been shown using blue bar and red bar, respectively. The abbreviations used in the panel are as follows; IMR-Intermediary metabolism and respiration, CHP-Conserved hypothetical protein, Vir- Virulence, detoxification and adaptation, IP- Information pathways, RP- Regulatory proteins, LM- Lipid metabolism, CW/CP- Cell wall and cell processes, ISP- Insertion sequences and phages, PE/PPE- Proline-glutamic acid (PE)/proline-proline-glutamic acid (PPE) sRNA – small RNA **(C)** The relative expression of differentially expressed transcripts was quantified after normalization to levels of housekeeping gene, *sigA*. The gene IDs are labelled on the x-axis and mean ± S.E. fold change obtained from three independent experiments is shown on y-axis.

### Generation of Rv0792c binding aptamer through SELEX

We next performed **S**ystematic **E**volution of **L**igands by **EX**ponential enrichment (SELEX) experiments to find DNA aptamers as a possible tool to identify epitope(s) which may bind small molecule inhibitors against Rv0792c. Thus, SELEX was performed using an 80-nucleotide long random ssDNA library. To diversify the sequences of aptamer library, SELEX binding experiments were performed using an error-prone Taq DNA polymerase (51). Prior to successive SELEX rounds, the double-stranded (ds) PCR products that were obtained were converted to single-stranded form, using previously reported methods (51). After 6 rounds of SELEX, the enrichment of Rv0792c-specific binders was determined by ALISA. As shown in Fig. 5A, the ssDNA pools from archived rounds (round 1 to round 6) were able to bind to Rv0792c protein. We observed saturation in the binding of aptamer pool to Rv0792c after round 3 of SELEX. Therefore, aptamer binders from round 6 of SELEX enrichment were cloned in the pTZ57R/T vector. The plasmid DNA from 17 randomly picked transformants was isolated and subjected to DNA sequencing. Next, the sequences of aptamer candidates were further analyzed using ClustalW and BioEdit software (Fig. 5B). Phylogenetic analysis revealed two preponderant groups among aptamer sequences (Fig. S3A). Group 1 is the largest and contains 11 aptamer candidates, while Group 2 included 6 aptamers (Fig. S3A). The base fraction analysis of these SELEX derived aptamer candidates evinced higher distribution of ‘G’ and ‘T’ residues over ‘A’ and ‘C’ (Fig. S3B). Based on primary sequence homology, 5 representative aptamer candidates were selected for further studies. Interestingly, sequence alignment studies revealed high homology between Rv0729c_1, Rv0729c_2 and Rv0729c_3 with the known DNA sequences for GntR family of bacterial transcription factors (Fig. S4, (13)).

**Figure 5:**
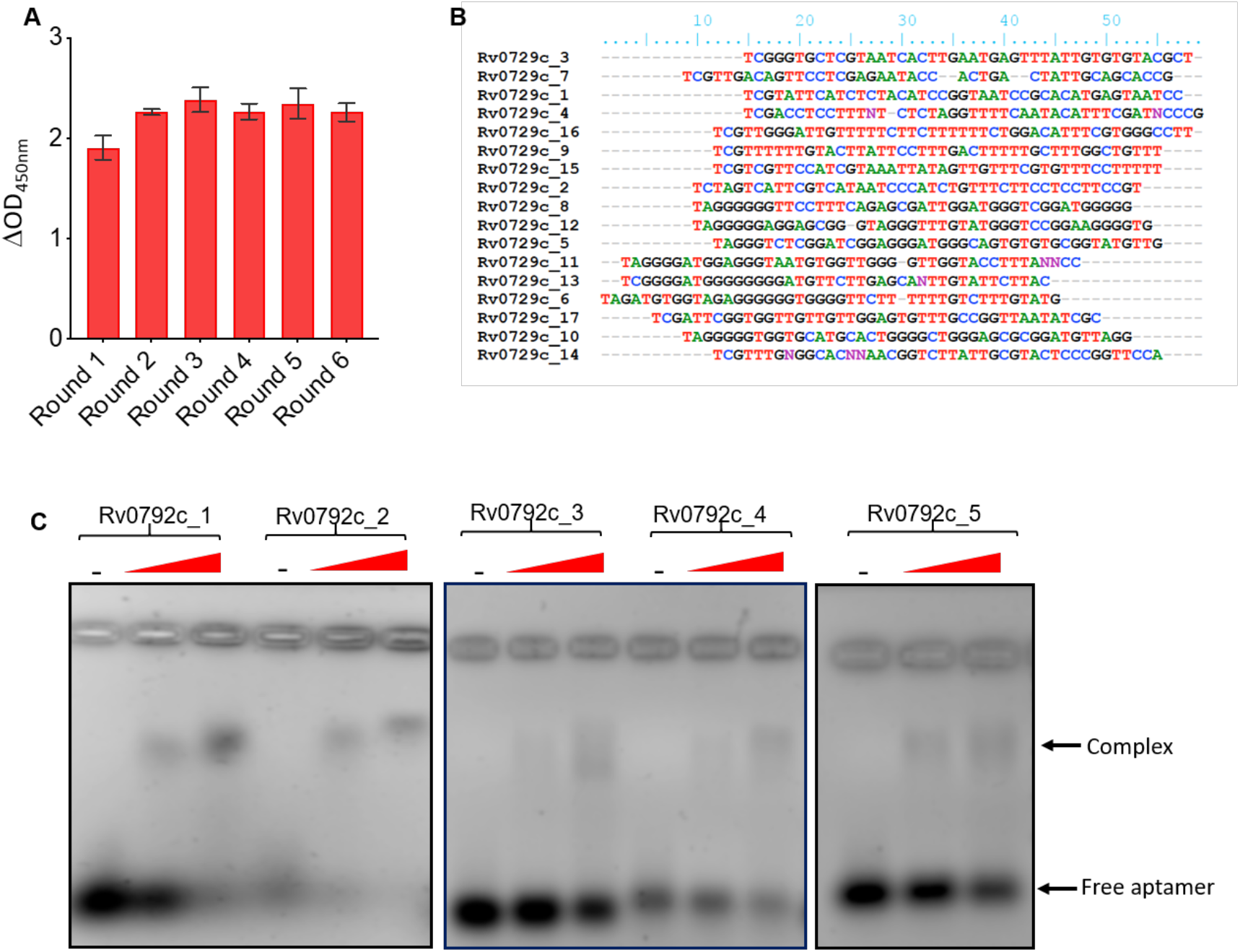
Selection and characterization of Rv0792c binding aptamers. **(A)** Round-wise enrichment of Rv0792c-binding aptamer pools (Round 1-6) assessed by ALISA. **(B)** Multiple sequence alignment of Rv0792c binding aptamer sequences. **(C)** Electrophoretic mobility Gel shift assay of selected aptamer candidates (100 pmol) with increasing concentrations of purified Rv0792c. The data shown in panel A is obtained from 2 replicates.

Next, to confirm the binding of aptamers with Rv0792c, EMSA assays using a panel of selected aptamer candidates (Rv0729c_1, 2, 3, 4 and 5) were performed. As shown in **Fig. 5C**, we observed that these aptamers interacted with Rv0792c at varying strength. Based on these findings, we selected three best aptamer candidates namely Rv0792c_1, Rv0792c_2 and Rv0792c_5 for further biochemical and functional characterization of Rv0792c. As shown in Fig. 6A, maximum binding with aptamers was observed with the wild-type protein, Rv0792c. Also, as expected, mutation of arginine 49 and glycine 80 abrogated the aptamer binding ability of Rv0792c. Based on these observations, we conclude that Rv0792c_2 displayed the highest binding for Rv0792c and in concordance with previous data, Arg49 and Gly80 are essential for binding of aptamers by Rv0792c (39, 40). In order to determine the role of Mg^2+^ ions in Rv0792c aptamer binding, ALISA assays were performed in the presence or absence of 10 mM EDTA. We observed that the inclusion of EDTA resulted in ∼90% reduction in aptamer binding to Rv0792c, thereby, indicating that Mg^2+^ ion is essential for aptamer-protein interaction (Fig. 6B). We determined the dissociation constant for binding of aptamer Rv0729c_2 with wild-type and mutant proteins and fitting of data was performed using the non-linear regression method. As expected, the highest binding was observed in wild-type protein with a Kd value of ∼51 nM. In comparison, Rv0792c^R49A^ and Rv0792c^G80D^ binds with Rv0729c_2 with Kd value of ∼ 74 nM and ∼ 120 nM, respectively (Fig. 6C). As Rv0792c belongs to the HutC family of GntR regulators, we next evaluated if L-histidine or urocanic acid (urocanic acid is an intermediate in the L-histidine catabolism) acts as effectors and alter the DNA binding ability of Rv0792c. Contrary to previous reports, we observed that the DNA binding ability of Rv0792c is not affected in the presence of L-histidine and urocanic acid (data not shown) (52–55). In order to identify Rv0792c effector molecules, ALISA was performed in the presence of various ligands. As shown in Fig. 6D, ALISA activity results indicate that Rv0792c aptamer binding ability did not increase in the presence of various ligands except L-arabinose. Approximately, 2.0-fold increase was seen in the presence of L-arabinose (Fig. 6D). These observations suggest that L-arabinose might act as an effector molecule for the DNA binding activity of Rv0792c or stabilizes the complex by maintaining the hydration layer.

**Figure 6:**
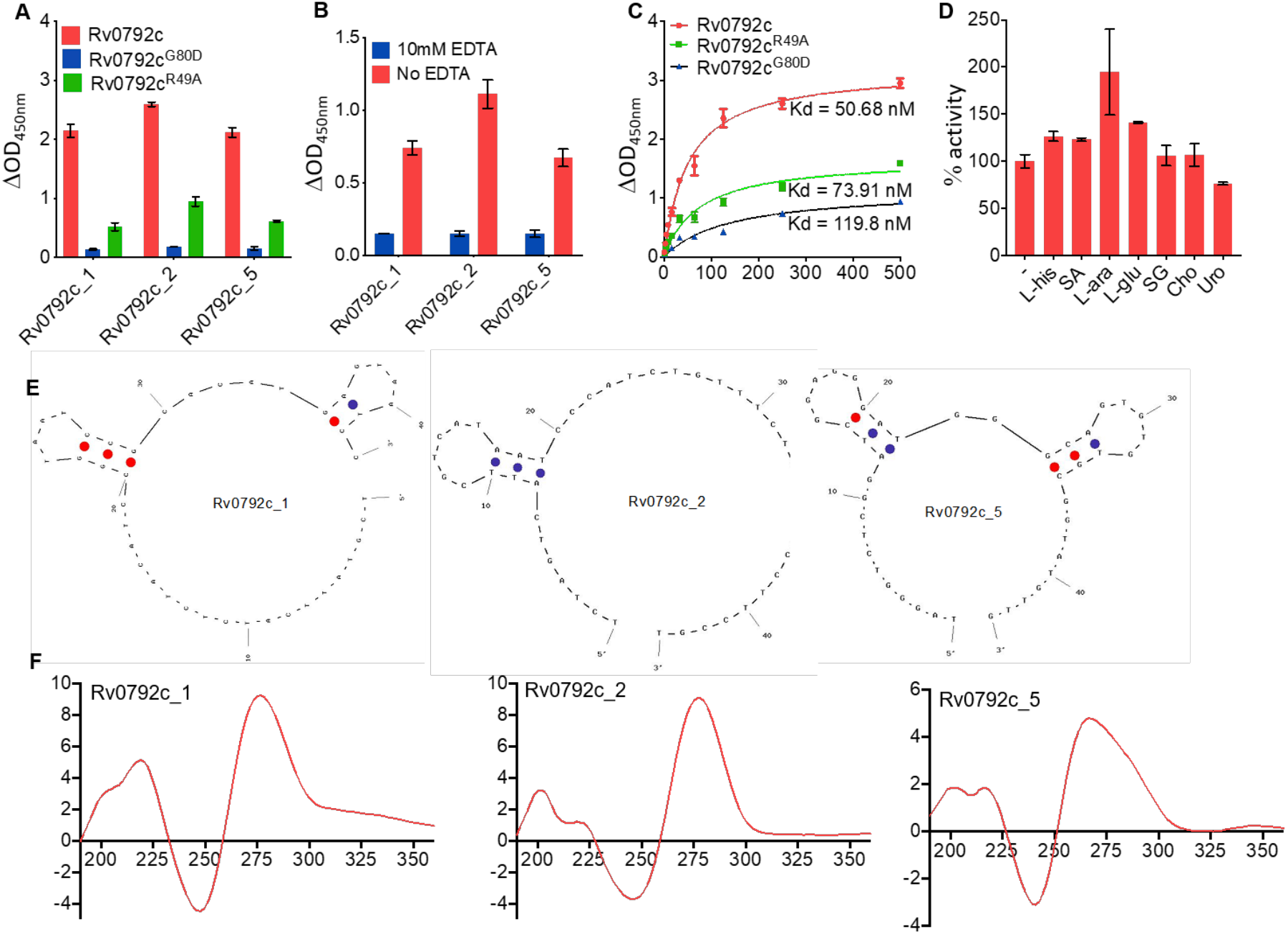
Binding and structural studies of selected aptamer candidates. (**A**) Binding of selected aptamer candidates (Rv0792c_1, Rv0792c_2 and Rv0792c_5) with wild type, Rv0792c^R49A^ and Rv0792c^G80D^. (**B**) Effect of presence of EDTA on the binding of Rv0792c_1, Rv0792c_2 and Rv0792c_5 aptamer candidates to Rv0792c protein. (**C**) Apparent dissociation constant curve derived through non-linear regression representing binding affinity (Kd) of Rv0792c_2 for wild type, Rv0792c^R49A^ and Rv0792c^G80D^. (**D**) Effect of various ligands on aptamer binding to Rv0792c protein. L-his, SA, L-ara, L-glu, SG, Cho and Uro represents L-histidine, stearic acid, L-arabinose, glucose, sodium glyoxylate, cholesterol and urocanic acid, respectively. (**E**) UNA fold predicated secondary structure of selected aptamer candidates. (**F**) Circular dichroism (CD) spectrum of Rv0792c_1, Rv0792c_2 and Rv0792c_5 aptamers. CD spectra indicate typical B-type stem-loop structure of aptamers. The data shown in panel a is obtained from 3 replicates. The data shown in panel b-d is obtained from 2 replicates.

### Secondary structure and circular dichroism spectral analysis

In order to determine the structures of the selected aptamers, they were subjected to UNA fold (https://eu.idtdna.com/UNAFold%3F). As shown in Fig. 6E, the secondary structure of selected aptamer candidates (Rv0792c_1, Rv0792c_2 and Rv0792c_5) showed a typical hairpin (stem-loop) like structure. Next, we performed circular dichroism (CD) studies for their conformational analysis. As shown in Fig. 6F, CD spectra of Rv0792c_1, Rv0792c_2 and Rv0792c_5 revealed a strong positive peak around 275-280 nm and a negative peak near to 240 nm. The observed spectra are in concordance with the spectra obtained for stem-loop like DNA (56, 57). However, the observed difference in the amplitude could be attributed to the variation in the sequence of these aptamers.

### SAXS Data Based Structural Model of Dimeric Rv0792c in the presence or absence of aptamers

We next acquired SAXS data to build a structural model in solution and identify the aptamer binding regions for Rv0792c. SAXS data was collected for Rv0792c at a concentration of 3.2 mg/ml as shown in Fig. S5A. The double logarithm mode of presentation confirmed a lack of aggregation or inter particulate effect in the protein sample (58). The inset in Figure S5A shows the Guinier region considering globular scattering nature and linear fit to the analysis confirming the monodisperse profile of the sample. Guinier analysis suggested that particle size to be characterized by a radius of gyration (R_g_) of about 3.3 nm (Table S3). Indirect Fourier transformation of the data provided frequency distribution of pairwise interatomic vectors which further provided an estimate of maximum linear dimension (D_max_) and R_g_ of 12.5 and 3.31 nm, respectively (Fig. S5B). Molecular mass estimation from different Bayesian models applied on experimental SAXS data suggested that the mass of the scattering particles was ∼ 63.2 ± 5.7 kDa supporting a dimeric state of association in solution (theoretical mass of monomer is 32.5 kDa).

As mentioned in materials and methods, a dummy residue model best representing the scattering shape of Rv0792c in solution was restored by averaging ten independent models and is presented in transparent map format with variation amongst models reflected as wire format (Fig. 7A and Fig. S6A). A normalized spatial disposition (NSD) value of 0.93 supported the similarity of the ten models solved and averaged for Rv0792c using SAXS data (Table S3). In order to compare the SAXS based envelope with structural model of Rv0792c, a sequence-based homology model was searched. The best sequence identity of 18.14% was observed between the 53-285 residues of Rv0792c with the solved structure for protein lin2111 from *Listeria innocua* Clip11262 (PDB deposition 3EDP; unpublished structure). We observed that most templates were similar in fold with a predicted association state of dimer. As stated in materials and methods, missing 52 and 16 residues from the amino and carboxy terminus, respectively, were modelled, their predominant conformation was oriented and subsequently attached to this central structural model of Rv0792c dimeric structure. Inertial axes of this structure was superimposed with those of SAXS based model for Rv0792c and similarities in the profile can be visually judged in the orthogonal views shown in Fig. 7A and Fig. S6A. Furthermore, a χ^2^ value of 1.3 between theoretical SAXS profile of the residue-level model of Rv0792c and experimental data supported a similarity between the two models in three-dimensions (Table S3). Zoomed-in image in Fig. 7A highlights that the two C-terminal extensions of chains bind each other, thus contributing to additional stabilization of the dimeric entity.

**Figure 7:**
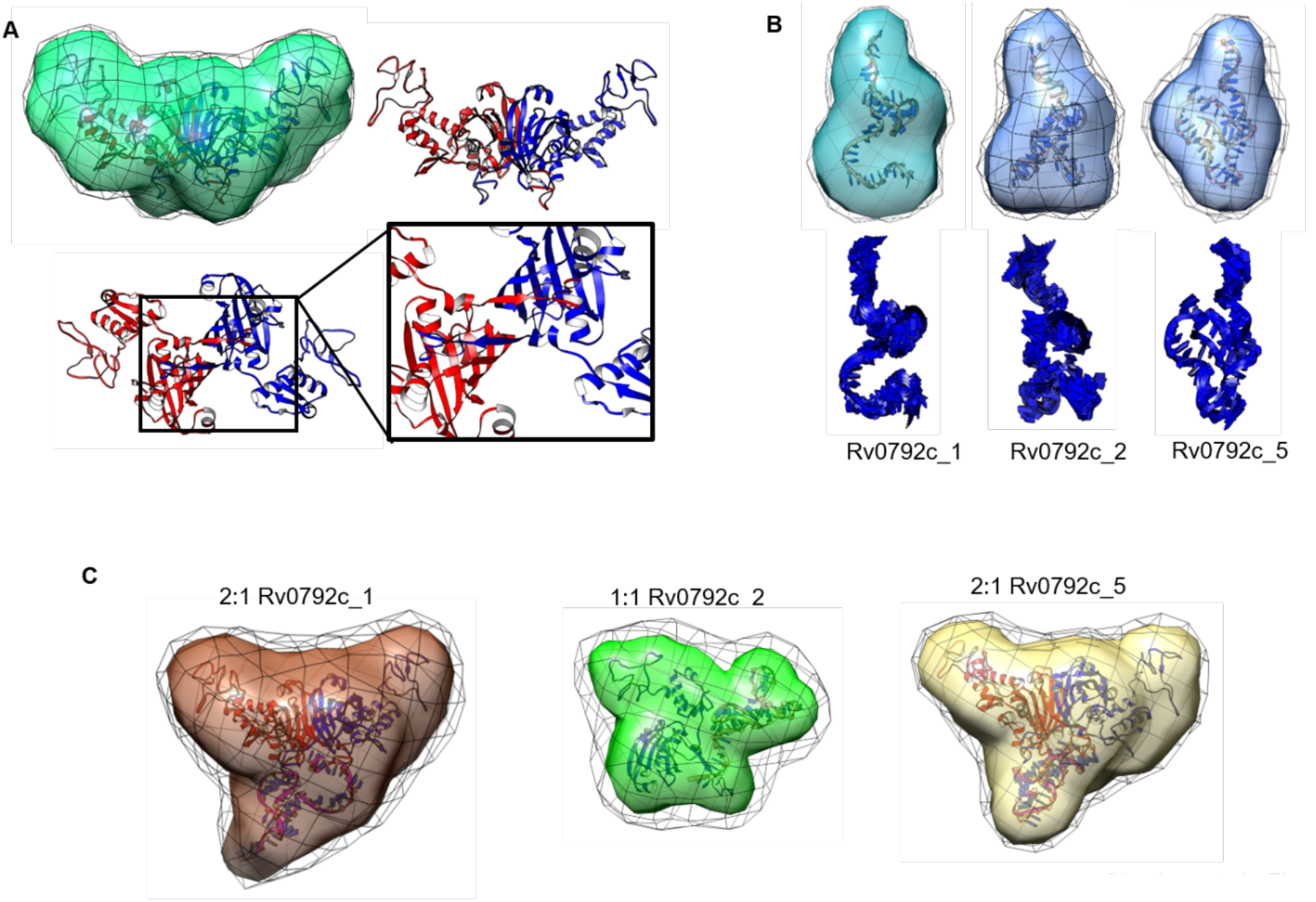
SAXS based structural models of unliganded protein Rv0792c, its binding aptamers and their complexes. (A) SAXS data-based envelope of Rv0792c dimer (green map) inertially aligned on structural model of protein (blue and red ribbon). The green map shows the common shape in ten solutions, and black mesh indicates the variation in them. Other panels show the C-terminal tail of each chain binds to the other stabilizing the dimeric association. (B) Upper panel shows the SAXS based dummy residue models of the unliganded aptamers Rv0792c_1, Rv0792c_2 and Rv0792c_5 as molecular map. Respective model is aligned with the inertial axes of its model from energy optimization (shown as ribbon). Lower panel shows the most collective normal mode frequency calculated for the energy optimized model of aptamer to reflect the inherent motion to the model. (C) SAXS data-based models of ternary and binary complexes of Rv0792c_1, Rv0792c_2 and Rv0792c_5 and protein Rv0792c are shown as molecular maps. The black mesh indicates the variations in the models solved for that complex. Inside each map, energy minimized model of the complex generated by SAXS data based selected result from docking of low energy conformation of aptamer on protein are shown as ribbons. Note that binding of Rv0792c_2 to protein leads to binary complex while other aptamers form stable ternary complexes.

Further, in order to perceive local and relative flexibility embedded in the computed structure of dimeric Rv0792c, low frequency normal modes accessible to the protein were calculated (Fig. S6B). The collective modes indicated that the N-terminal domain moved in synchronized mode independent of the central β-barrel type dimeric contact. The C-terminal tail of the proteins also move up and down the interacting β-barrel and linker connecting the barrel and N-terminal domain of the other chain in dimer. These theoretical analyses imply that the C-terminal ends of the dimer remain attached to each other chain. Next, using SAXS data analysis, solution shape parameters, association state, and structure of Rv0792c binding aptamers were determined in their unliganded state (Fig. 7B). Double Log profile of SAXS data from aptamers confirmed lack of any aggregation or interparticulate effect in the samples (Fig. S5C). Guinier analysis for globular scattering profiles are shown as inset and linearity of the fits in low q range further validated monodisperse nature of aptamers. The parameters deduced for predominant scattering shape of aptamers are listed in Table S3. In summary, aptamers had R_g_ and D_max_ in range of 1.8-2.1 nm and 7.1-8.6 nm, respectively. For all aptamers, the calculated P(r) indicated a “tailing” at higher r values suggesting flexible ends about core shape (Fig. S5D). Using SAXS data profiles, their estimated molecular masses were in the range of 12.9 – 13.5 kDa, clearly supporting a monomeric state of these ssDNA molecules in solution. Their dummy residue models solved within SAXS data-based constraints are shown in Fig. 7B. As mentioned in methods, considering monomeric status, predominant low energy conformations of the ssDNA aptamers were calculated in implicit dielectric of 80 (representing water), and the best-resembling conformation was selected using lowest χ^2^ value between the calculated SAXS profile for the conformation and experimental data. These models for aptamers are shown overlaid on SAXS data-based model and alone in Fig. 7B. The additional views are shown in Figs. S7A, S7B and S7C. Relative to Rv0792c, NSD values for aptamers in the range of 0.5-0.7 indicated differential nature of ten models solved for the three aptamers (Table S3). This implied relatively higher inherent disorder in the unliganded aptamers as monomers. Similar disorders were observed in the higher terminal motions in the residue level models computed for these aptamers. Pertinently, it also explained the extended nature of their computed P(r) curves.

Having characterized that Rv0792c protein adopts dimeric state in solution, and all the binding aptamers are monomers, next set of SAXS data was acquired on molar mixtures of the protein and individual aptamers (ratio was computed for dimeric to monomeric state of Rv0792c and ssDNA) (Fig. S5E and S5F). It is important to state here that concentration of the molecules were higher than the estimated binding constant of the protein and DNA molecules, supporting a higher order of binding between available molecules and scope of none or little unliganded molecules in samples used for SAXS data collection. Double Log plot and Guinier analysis of the Rv0792c_1, Rv0792c_2 and Rv0792c_5 aptamers support that the scattering molecules did not aggregate or underwent inter particulate effect upon mixing (Fig S5E). While mixtures of Rv0792c with aptamer Rv0792c_5 and Rv0792c_1 showed D_max_ and R_g_ values of 12.5 and 3.4-3.5 nm, respectively, the complex of GntR and Rv0792c_2 aptamer adopted a decreased D_max_ and R_g_ of 9.8 and 3.1 nm, respectively (Table S3). The lower dimensions of complex with Rv0792c_2 was correlated with SAXS based molecular mass prediction which indicated mass of ∼ 44 kDa for this complex, and ∼75 kDa for samples with Rv0792c and aptamers Rv0792c_5 and Rv0792c_1. This result indicated that Rv0792c_5 and Rv0792c_1 binds to dimeric protein and does not alter the association state of Rv0792c into monomers. In contrast, binding of Rv0792c_2 aptamer induces dissociation of dimeric proteins into monomers. The observed differences with R_g_, D_max_ values and P(r) profiles between these Rv0792c-aptamer complex, unliganded Rv0792c and aptamers supported that protein and aptamer molecules were bound to each other during data collection. Shapes restored for these scattering species showed higher NSD values than unliganded aptamers.

As mentioned in materials and methods, results from SAXS data analysis revealed that Rv0792c_5 and Rv0792c_1 form 2:1 complex with Rv0792c, but binding of Rv0792c_2 dissociates Rv0792c dimer into monomers. Accordingly, the low energy structures of aptamers were docked on Rv0792c dimers for Rv0792c_5 and Rv0792c_1 aptamer, and on monomer for Rv0792c_2 aptamer to obtain models for their complexes. For latter, approximation was made that no large shape change occurred theoretically detaching monomer from dimer of Rv0792c. Different poses of docked aptamers on Rv0792c were filtered to correlate with the shape solved for the complexes (Fig. 7C and Fig. S8). The models selected for Rv0792c: aptamer complexes indicated that aptamers bind to the C-terminal portion of Rv0792c (Fig. 8A). Energy minimization of the residue-level models of complexes obtained from docking indicated that while Rv0792c_5 and Rv0792c_1 aptamer coalesced with dimer interface, Rv0792c_2 aptamer induced opening of the C-tail latch of Rv0792c. Probably, this last event in case of Rv0792c_2 weakens the protein-protein interaction between Rv0792c and leads to eventual formation of 1:1 complex. In summary, all aptamers remain monomer in the presence or absence of Rv0792c and bind to its C-tail region, and some interaction extends to the stretch encompassing PRG (residues 40 – 42) of Rv0792c protein (boxed in the zoomed image in lower panel in Fig. 8A).

**Figure 8:**
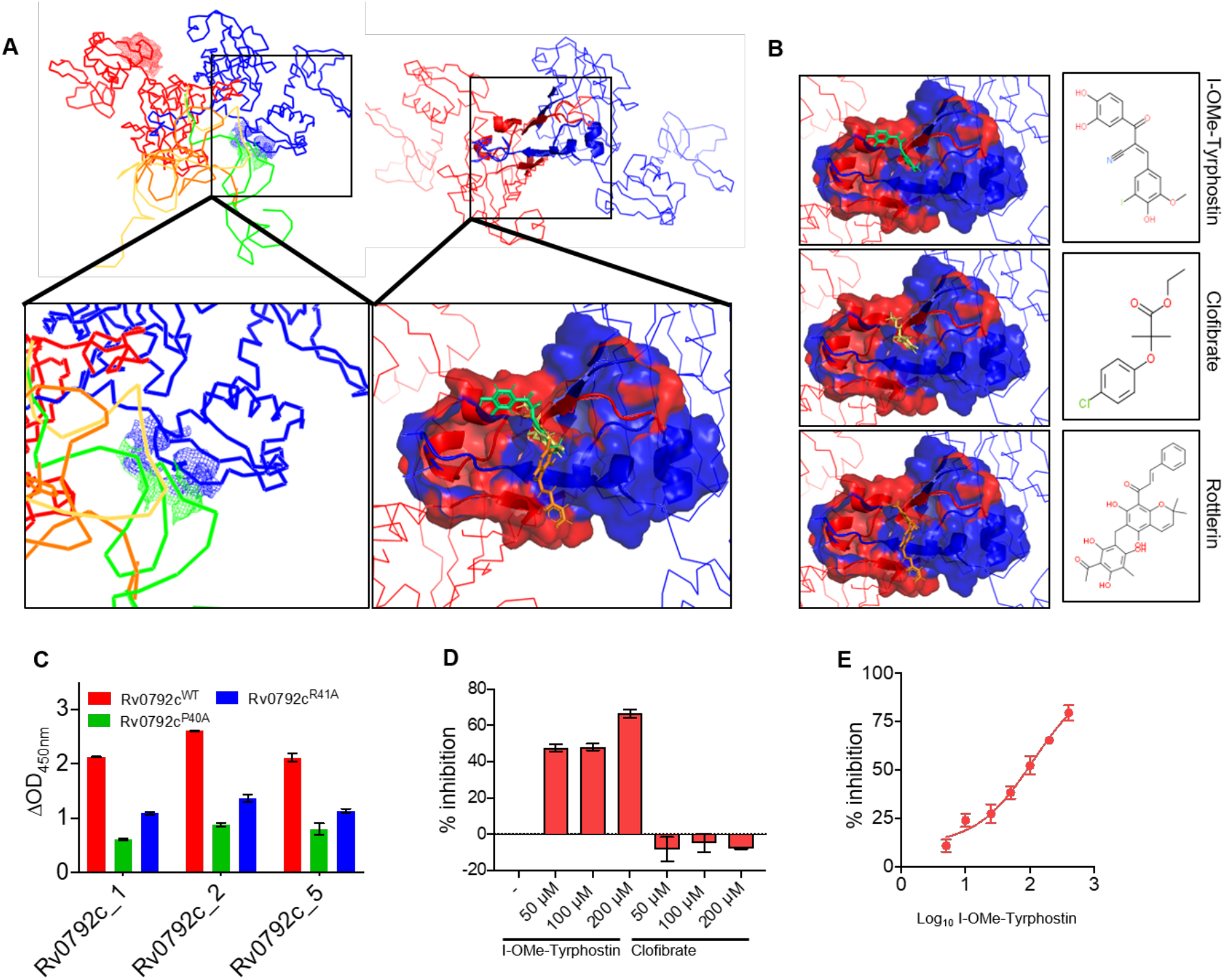
Identification of I-OMe-Tyrphostin as a small molecule inhibitor of Rv0792c. Left upper panels show ribbon representation of (left) the similar binding of the three aptamers in space near the C-terminal dimerization region, and (right) shows the segment which is common to binding of all aptamers. The zoomed in panel in lower left part show how aptamers binding to C-terminal part extend interaction to the functionally critical PRG motif in one chain of protein (shown as blue mesh). The zoomed in image in the right lower panel shows the docking of three best hits on the surface common to binding of aptamers. The receptor is shown in surface map mode with colouring of the chain of the dimer. The molecules are shown as sticks. (B) The right three panels show the docked pose of individual molecules with their chemical structures in inset. From top to bottom, hit molecules are: (2E)-2-(3,4-Dihydroxybenzoyl)-3-(4-hydroxy-3-iodo-5-methoxy phenyl) acrylonitrile (*I-OMe- Tyrphostin);* 2-(4-Chlorophenoxy)-2-methylpropionic acid ethyl ester (*Clofibrate*); and (2E)- 1-[6-(3-Acetyl-2,4,6-trihydroxy-5-methylbenzyl)-5,7-dihydroxy-2,2-dimethyl-2H-chromen- 8-yl]-3-phenyl-2-propen-1-one (*Rottlerin*). (C) Binding of selected aptamer candidates (Rv0792c_1, Rv0792c_2 and Rv0792c_5) with wild type, Rv0792c^P40A^ and Rv0792c^R41A^. (D) The aptamer binding activity of Rv0792c was determined by ALISA in the presence of I-OMe- Tyrphostin and Clofibrate at 50 μM, 100 μM and 200 μM concentration. (E) The binding of Rv0792c with aptamer was performed in the presence of different concentration of I-OMe- Tyrphostin as described in Materials and Methods. The data shown in panel c-e is obtained from 2 replicates.

### Computational docking to identify small molecule inhibitors for Rv0792c

As seen from shape restoration and docking data, all screened aptamers were binding to C-terminal dimerizing segment of Rv0792c protein. Additionally, binding of Rv0792c_2 induces dissociation of Rv0792c dimer. Presuming that this segment is key to structural organization of Rv0792c and small molecules capable of binding this segment may alter the native functioning of this protein, we used the aptamer binding segments to screen for molecules which can bind to this protein. Fig. 8B shows the defined aptamer binding region and three best hits in their lowest energy pose with the receptor. These molecules were I-OMe-Tyrphostin, Clofibrate and Rottlerin in the decreasing order of their relative docking score. Next, we performed ALISA experiments to investigate whether (i) PRG motif is required for aptamer binding to Rv0792c and (ii) whether the small molecules identified from computational docking can inhibit the binding of Rv0792c_2 aptamer to Rv0792c. In order to determine the role of PRG motif in binding to these aptamers, Rv0792c harboring Pro^40^-Ala^40^ and Arg^41^-Ala^41^ mutation was cloned, expressed and purified as (His)_6_ tagged protein. In concordance with SAXS based modeling results, we observed that mutation of either proline 40 or arginine 41 to alanine abrogated the aptamer binding ability of Rv0792c (Fig. 8C). Of the three compounds, we were able to procure only 2 compounds with assured purity. Both compounds did not exhibit solubility issues and were evaluated for their ability to inhibit Rv0792c enzymatic activity. As shown in Fig. 8D, among these two compounds, I-OMe-Tyrphostin was able to inhibit the binding of aptamer to Rv0792c protein by ∼60% at 200 μM concentration. We noticed no inhibition of aptamer binding in the presence of Clofibrate even at 200 μM concentration (Fig. 8D). We also observed that I-OMe-Tyrphostin was able to inhibit Rv0792c activity in a concentration dependent manner. As shown, the small molecule inhibited aptamer binding with an IC_50_ of ∼ 109 nM (Fig. 8E). Taken together, this is the first study, where we show that GntR homolog, Rv0792c from *M. tuberculosis* is essential to establish infection in host tissues. We also report novel aptamers which bind to the dimerizing segment of the protein, and used this information to identify an FDA-approved drug which can conceptually act as an inhibitor of Rv0792c protein.

## Discussion

Transcriptional regulation has been shown to be essential for adaptation of various bacterial pathogens upon exposure to unfavourable and harsh environmental conditions. *M. tuberculosis* is a highly successful intracellular pathogen owing to its ability to sense external stimulus, reprogram its transcription machinery and persist in host tissues. The *M. tuberculosis* genome encodes for several transcription regulators that have been demonstrated to be essential for its growth *in vitro* or *in vivo*. GntR family of transcription factors are widespread among prokaryotes and are involved in various processes such as (i) carbon metabolism, (ii) motility, (iii) antibiotic production, (iv) drug tolerance, (v) biofilm, (vi) virulence and pathogenesis (15–18, 23). GntR is a relatively new and still poorly characterised family of transcriptional regulator. Despite the presence of GntR homologs in the genome of *M. tuberculosis* their biological functions have not been well established extensively (24–26). In the present study, we have used different approaches to characterize Rv0792c protein (HutC homolog) from *M. tuberculosis*.

GntR family of transcription regulators binds DNA as dimers and subsequently either repress or activate their own transcription (13). Using sedimentation ultracentrifugation and SAXS data analysis, we show unambiguously that Rv0792c exists as dimer in solution as reported for other GntR homologs (59). Although the sequence similarity between amino-terminus DNA binding domains is 25%, the residues essential for interaction with DNA are highly conserved (10). Sequence alignment studies revealed that the highly conserved helix-turn-helix DNA binding domain of Rv0792c spans amino acid between 21-86 residues. As expected, substitution of Glycine at position 80 to aspartic acid abrogated the DNA binding properties of Rv0792c in EMSA and ALISA-based assays. These results implicates that glycine residue is essential for DNA binding properties as reported in the case of *E. coli* FadR (41). In GntR family, the conformation of DNA binding motif is altered by binding of effector molecules. The binding of fatty acyl-CoA negatively regulates DNA binding activity of FadR in *E. coli* (60). Similarly, DNA binding of AraR in *B*. *subtilis* is repressed by binding of arabinose to the effector binding domain (61). The activity of HutC subfamily has also been reported to be regulated by N-acetyl-glucosamine and urocanic acid (55, 62). We also evaluated the ability of various effectors to regulate DNA binding activity of Rv0792c and observed that inclusion of L-histidine and urocanic acid couldn’t affect the DNA binding ability of Rv0792c. However, arabinose increased the DNA binding ability of Rv0792c by ∼2.0-fold which suggests that L-arabinose might be the effector molecule for Rv0792c. Nevertheless, regulation of Rv0792c might still be fine-tuned by this specific ligand interaction and/or by some unknown ligand that needs to be investigated in near future.

In order to investigate GntR role in physiology, stress adaptation and virulence, Rv0792c mutant strain was generated using temperature sensitive mycobacteriophages. We noticed that the colony morphology, growth pattern and biofilm formation of the mutant and parental strain were comparable. Transcriptional regulators are well known in mediating mycobacterial stress adaptation. In order to mimic the environmental clues as encountered by mycobacterium within the host macrophage or granuloma, the survival of various strains was evaluated in different stress conditions. The mutant strain was compromised for survival upon exposure to oxidative stress and cell wall damage. However, no differences were observed between wild-type and mutant strains upon exposure to other stress conditions such as nitrosative, low oxygen, nutrient starvation and acidic pH. The deletion of Rv0792c in the *M. tuberculosis* genome also didn’t alter the survival upon exposure to various drugs with a different mechanism of action. GntR family of transcription regulators have also been shown to be involved in the virulence of other bacterial pathogens (63–66). Notably, we observed that the Rv0792c mutant strain was markedly defective in establishing infection in lungs and spleens of guinea pigs. Both 4 and 8-week data showed significantly higher lungs and splenic bacillary load of guinea pigs infected with the wild type strain in comparison to the mutant strain. In agreement, less tissue damage was seen in H&E stained sections from guinea pigs infected with the mutant strain. The granuloma formation was seen in sections from wild type (necrotic) and complemented strains (non-necrotic). These observations implicate that Rv0792c plays a key role in *M. tuberculosis* virulence and is essential for establishing infection in host tissues. RNA-seq analysis constitutes an important functional framework to determine which gene or a subset of genes response impacts survival mechanisms deployed during host infection. To further understand the mechanisms associated with the attenuation of Rv0792c mutant strain in host tissues, transcriptomic profiles of wild-type and mutant strain were compared. This analysis revealed several important genes linked to M. *tuberculosis* virulence and survival such as *leuC* (Rv2987c), *desA2* (Rv1094), *efpA* (Rv2846c), *clgR* (Rv2745c) were differentially expressed. The transcript of several genes belonging to either PE/PPE or toxin-antitoxin modules or lipid metabolism were also downregulated in the mutant strains. We also observed that the transcript levels of Rv0793 (gene neighbouring to Rv0792c) and *mymA* operon (shown to be upregulated in acidic conditions) were increased in the mutant strain (50, 67). These findings clearly suggest that Rv0792c regulates the expression of “subset” of genes that enables the bacteria to adapt and persist in host tissues.

SELEX strategy was employed to search for novel ssDNA aptamers capable of tightly binding Rv0792c. Extended aim was to utilize the aptamer binding information to screen for small molecules which may efficiently bind Rv0792c and act as inhibitors of its function. The direct interactions between Rv0792c and SELEX derived DNA aptamers were confirmed using EMSA and ALISA. Among the identified, three DNA aptamer candidates (Rv0792c_1, Rv0792c_2 and Rv0792c_5) showed good binding to Rv0792c albeit with varied intensity. This difference in the intensity of the DNA-protein complex evinced the differential rate of aptamer–target complex association and dissociation (56, 68). Interestingly, the binding of all three aptamers was abrogated by the substitution of glycine 80 to aspartic acid and arginine 49 to alanine in the helix-turn-helix motif. This is possibly because of two reasons: (i) aptamer selection was performed with the wild-type protein and (ii) these residues (glycine 80 and arginine 49) play a critical role in maintaining the protein structure where aptamer binds (41). Notably, the presence of EDTA also abrogated the aptamer binding to Rv0792c indicating that Mg^2+^ is essential to maintain the active conformation of aptamers required for protein binding. This finding is in agreement with the previous reports where the role of divalent ion in aptamer binding to cognate protein target has already been established (69). Interestingly, all these aptamer candidates displayed Kd in the nanomolar range, an observation which is in concordance with the previously reported protein binding aptamers (29, 70, 71). Notably, the primary sequence of Rv0729c_1, Rv0729c_2 and Rv0729c_3 DNA aptamer candidates designed against the Rv0792c, a HutC protein showed high similarity to DNA binding sequences of FadR subfamily transcription factors (13). We observed that the level of similarity was much higher in the case of Rv0729c_1, Rv0729c_2 compared to Rv0729c_5. This pattern clearly indicates the possible role of these nucleotides to provide affinity to bind the transcription factor of the GntR family. *In silico* structure prediction and their validation by CD clearly demonstrates the presence of stem-loop like structures which are very common among protein binding DNA aptamers (70, 72). As mutation of arginine and glycine abrogated aptamer binding, our data suggests that aptamers are binding to a region in Rv0792c that is essential for DNA binding. Therefore, we hypothesized the epitope at which these aptamers bind may be functionally important and a small molecule binding to this site may impede the functional profile of the protein.

Further, to gain insight into the complexes of aptamers bound to Rv0792c, SAXS data analysis and molecular modelling was utilized. Analysis of the unliganded protein and aptamers showed that in solution their association state is predominantly dimer and monomer, respectively. The dimeric status of Rv0792c correlated well with the AUC data, which showed presence of minor higher-order associated species too. Interestingly, mixing of aptamers to dimeric Rv0792c showed that while one molecule of Rv0792c_1 aptamer and Rv0792c_5 aptamer binds to one dimer of Rv0792c, binding of Rv0792c_2 aptamer led to dissociation of the Rv0792c dimer into monomer. *In silico* molecular modelling steered and selected within constraints from experimental SAXS data provided a key insight that Rv0792c dimerizes across its C-terminal, and the extended C-tail wraps around each other chain that provides additional stability to the dimer. The dimeric status or even association architecture is not novel to the GntR family of proteins, but the unique wrapping up of C-tail on each other chain opens up queries on its functional relevance and possible uniqueness in this family of regulators. Solution scattering data supported that the aptamers remained predominantly monomer. Interestingly, the structural analysis revealed that the exposed side of the dimeric Rv0792c is also the interaction site of the three aptamers that were identified from the SELEX study. Taken together, SAXS data provided insight that binding of Rv0792c_2 aptamer induces rearrangement(s), which leads to dissociation of the dimer of Rv0792c protein.

Taking cue from the poses of aptamers on Rv0792c dimer, we considered using the interacting residues in the protein to screen for small molecules which may even compete with the binding of aptamers. The two molecules from the identified top hits were experimentally evaluated in our aptamer binding assays and we observed that I-OMe- Tyrphostin was able to inhibit binding Rv0792c_2 aptamer to Rv0792c. It is worth mentioning here that Rv0792c_2 binds with the highest affinity to Rv0792c, so it can be safely extrapolated that Tyrphostin analog may also competitively inhibit binding of other two aptamers. This molecule, I-OMe-Tyrphostin and its analogs have been assayed before for their potential as epigenetic regulator (73, 74) No specific correlations have been made with *M. tuberculosis*, except a recent study which identified new inhibitors for the Pup proteasome system in *M. tuberculosis* (https://doi.org/10.1101/796359). They report that I-OMe- Tyrphostin and Tyrphostin inhibit Dop, a depupylase from *M. tuberculosis*. The future course of our experiments will explore the efficacy of this small drug molecule in inhibiting the growth or survivability of *M. tuberculosis* in different assays or models. Definitely, being an approved drug, any efficacy against *M. tuberculosis* will enable its quick translation. In conclusion, we have (i) delineated the role and contribution of GntR-like factors in *Mtb* physiology, stress tolerance and pathogenesis and (ii) also identified small molecule inhibitor against Rv0792c, an *in vivo* essential transcription factor.

## Supplementary Information

**Figure S1:**
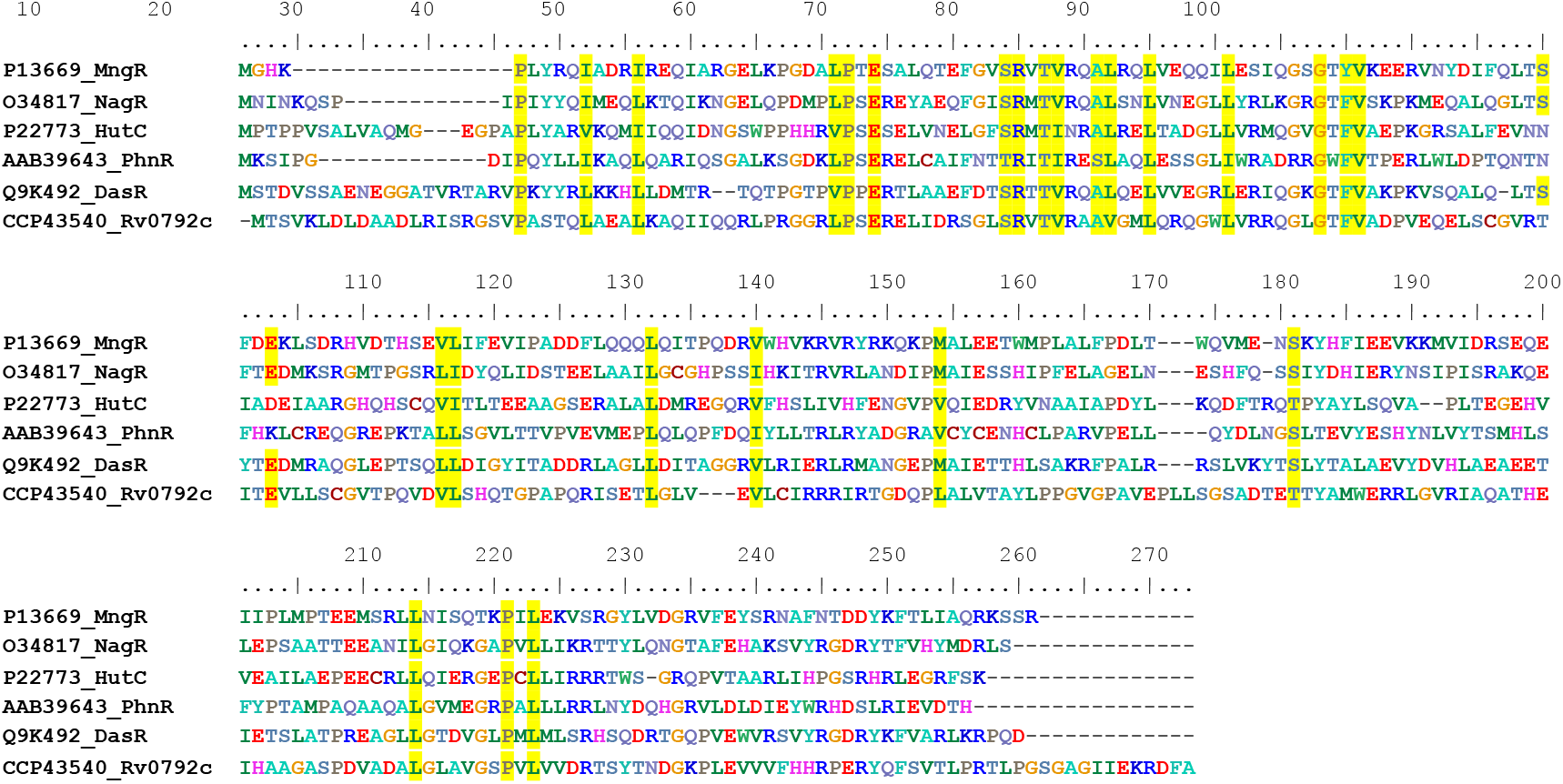
Multiple sequence alignment of putative Rv0792c homologs from various bacterial pathogens. Multiple sequence alignments among different proteins were prepared using Clustal W and formatted using BioEdit sequence alignment editor. The conserved residues among different proteins are highlighted in red. The proteins used in alignment are; CCP43540 (Rv0792c, *M. tuberculosis*), P13669 (MngR, *E. coli*), O34817 (NagR, *B. subtilis*), P22773 (HutC, *P. putida*), AAB39463 (PhnR, *S. enterica*) and Q9K492 (DasR, *S. coelicolor*).

**Figure S2:**
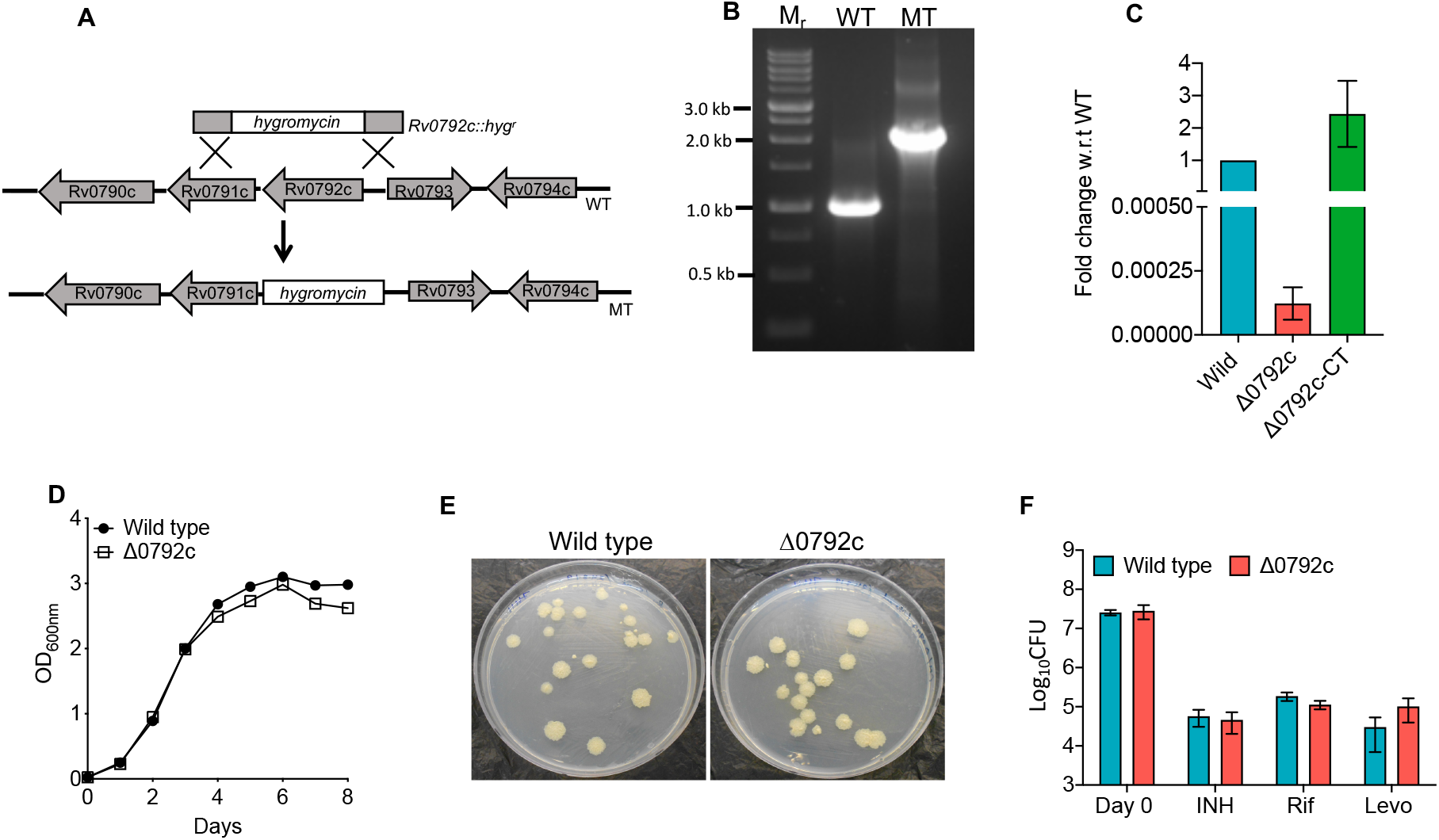
**(A) Schematic representation of Rv0792c locus in wild type and mutant strain of *M. tuberculosis.*** The gene for Rv0792c was replaced with the hygromycin resistance cassette in the mutant strain by homologous recombination using temperature sensitive mycobacteriophages **(B-F) Characterization of Rv0792c mutant strain of *M. tuberculosis***. The disruption of Rv0792c in the mutant strain was confirmed by PCR (B) and qPCR analysis (C) using gene locus specific primers. (D) The growth kinetics of wild type and Rv0792c mutant strain was compared by measuring the absorbance at 600nm. (E) The colony morphology of wild type vs mutant strain is shown in this panel. (F) For *in vitro* drug susceptibility assays, mid-log phase cultures of various strains were exposed to various TB drugs. The data shown in this panel is mean ± S.E. of results obtained from two independent experiments.

**Figure S3:**
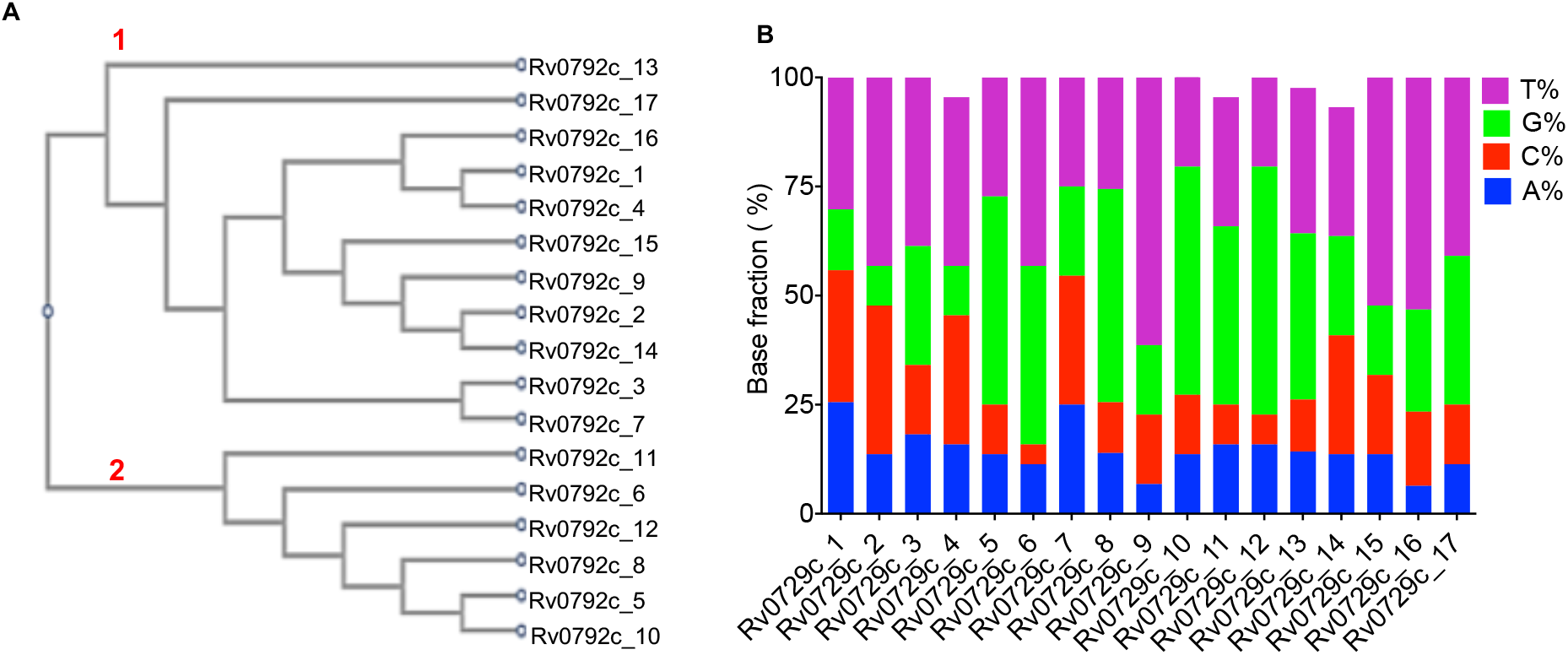
**(A)** Phylogenetic tree analysis of aptamer sequences using online tool ClustalW. The numbers in red denote the preponderant groups in the phylogenetic tree. **(B)** Base fraction analysis of aptamer candidates. We observed that the majority of selected aptamers evinced bias towards ‘T’ richness.

**Figure S4:**
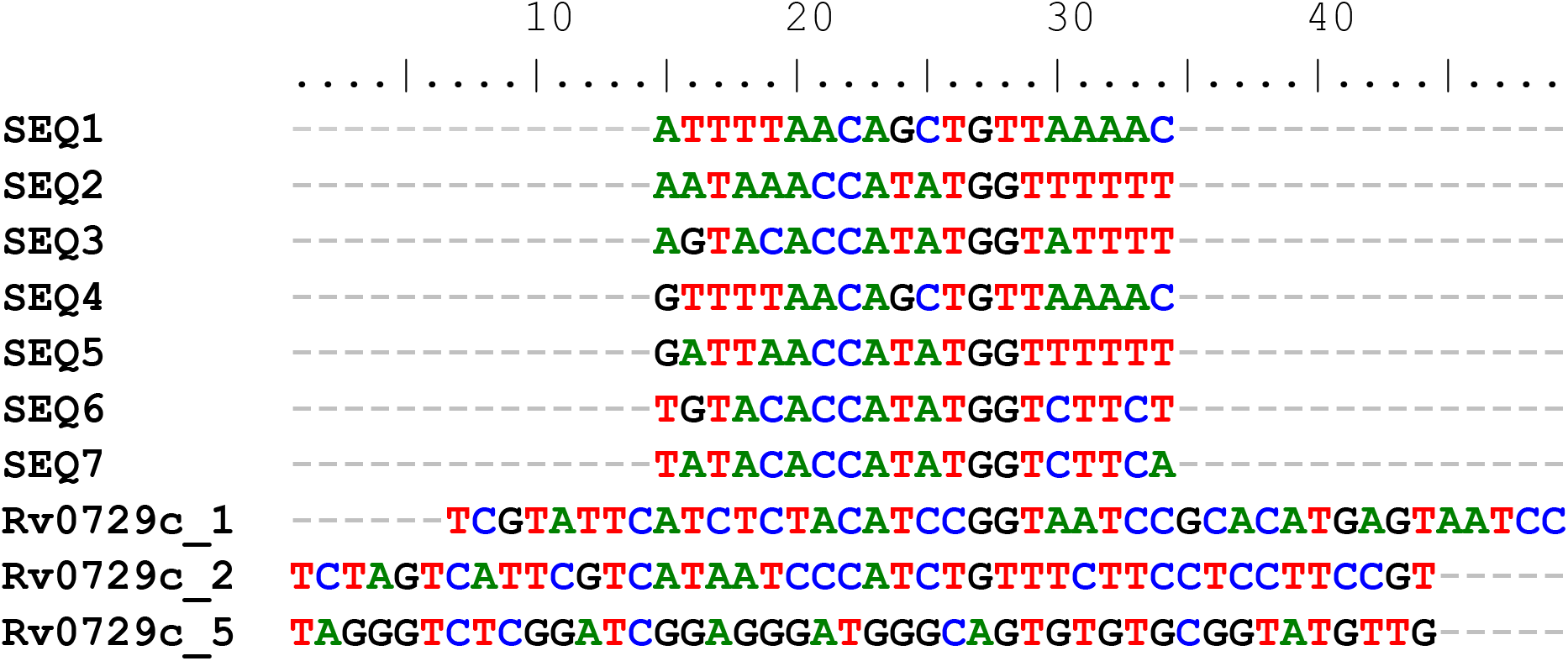
Multiple sequence alignment of Rv0792c_1, Rv0792c_2 and Rv0792c_5 aptamers with the known DNA sequences having affinity for transcription factors of GntR family. Seq1 to Seq7 represent the known DNA sequences.

**Figure S5:**
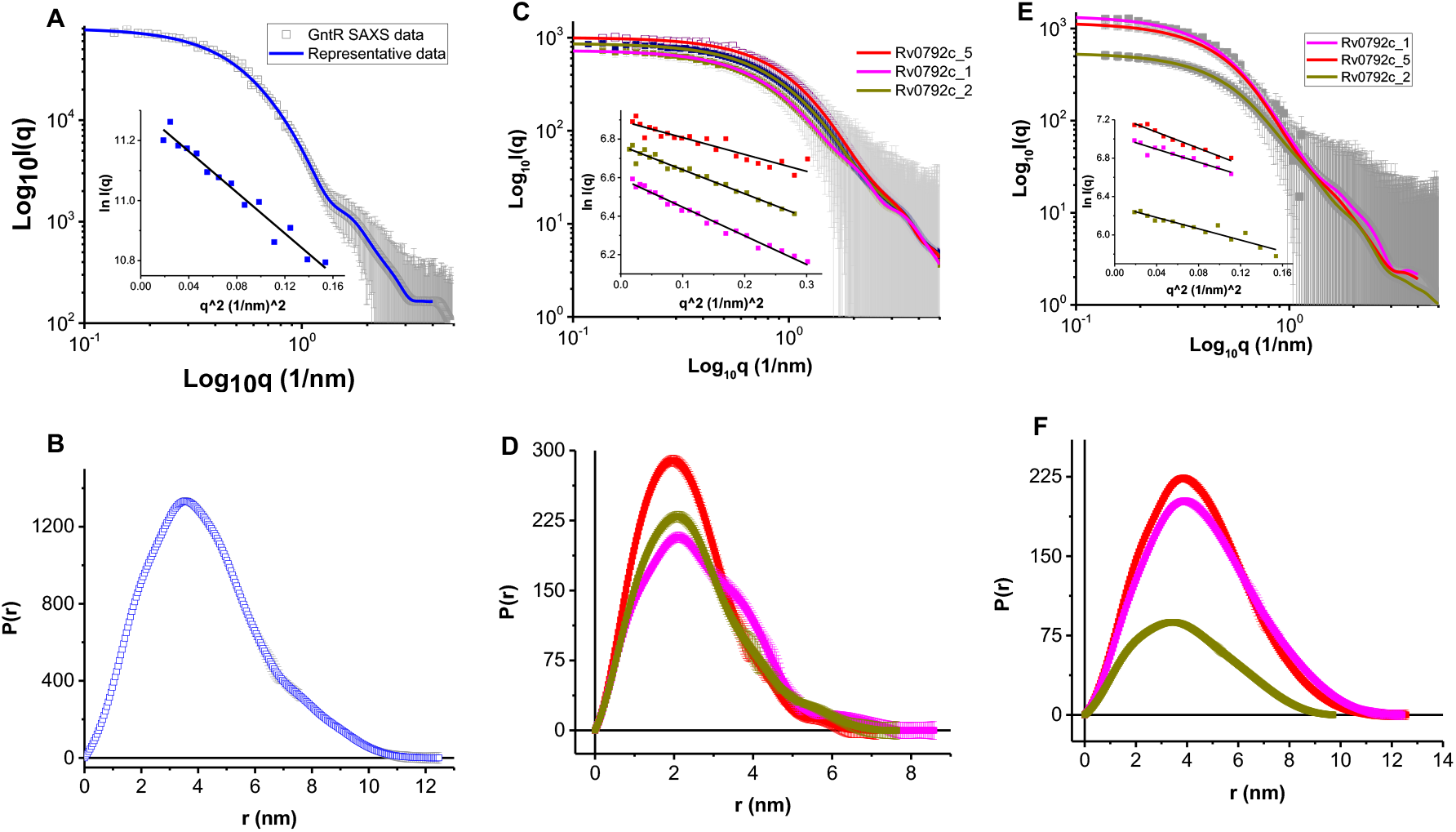
The panel **(A)** shows SAXS data acquired from a sample of Rv0792c at concentration close to 3.2 mg/ml (grey square symbols). Inset shows the linear region of the SAXS dataset in Guinier analysis considering globular scattering profile. **(B)** This panel shows the distance distribution profile of the interatomic vectors inside SAXS profile of the protein, and the blue line in the left panel plot shows the SAXS profile of the estimate. **(C-F)** Same analyses of the unliganded aptamers (**C, D**) and 1:1 molar mixture of protein and aptamers (**E, F**) are shown, respectively.

**Figure S6:**
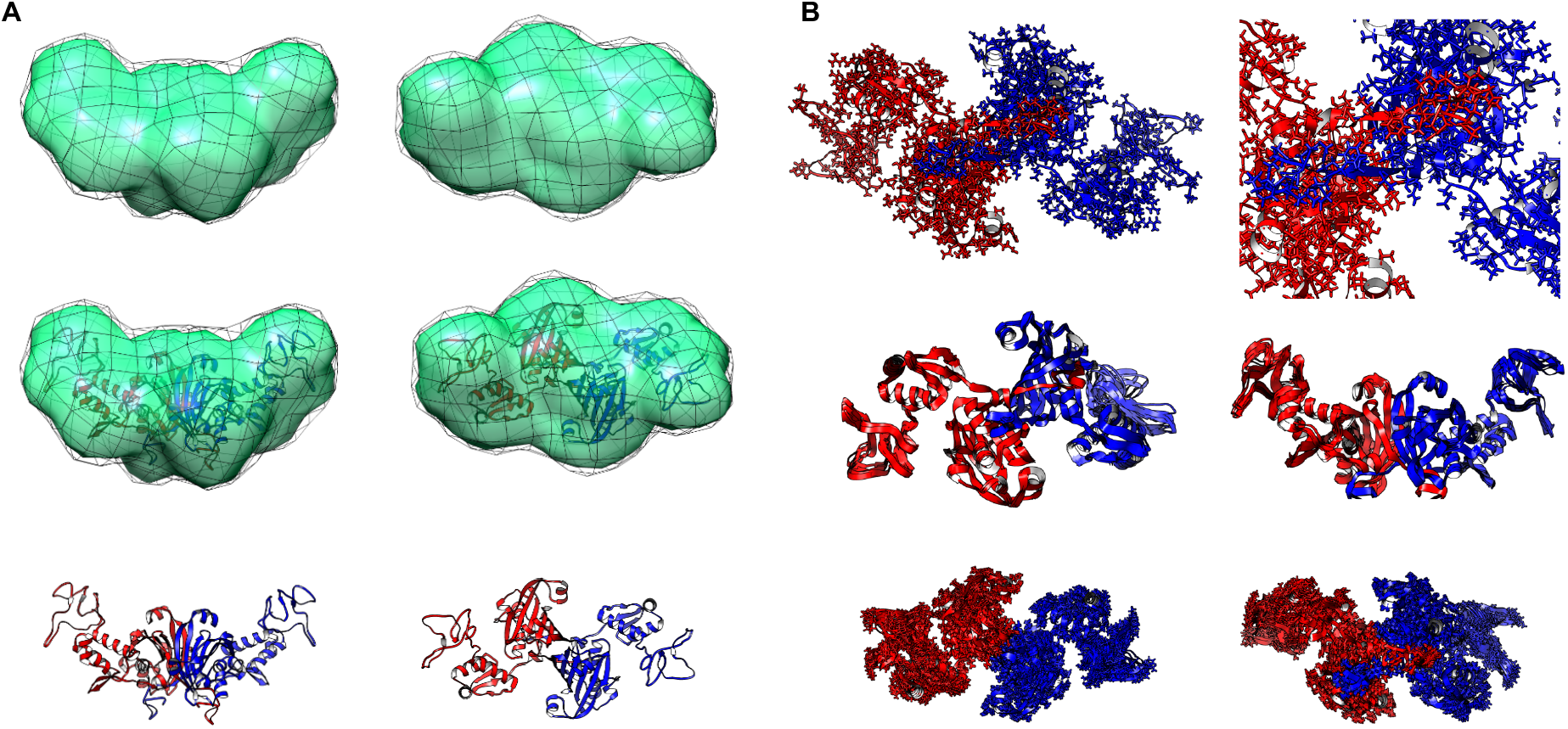
**(A)** Two orthogonal views compare the SAXS data based scattering envelope shape of Rv0792c and its structure solved by a combination of homology modelling and energy minimization. The green map represents the shape profile which was calculated to be common in ten dummy residue models of the protein. The variation is depicted as black mesh. The bottom panels show rotated views of the residue detail model of protein which was found to be a dimer and is shown in ribbon format. The central panel shows the inertial axes overlay of the two models indicating their similarity in three-dimensional space. **(B)** The calculated low frequency normal modes of motion accessible to the model of the Rv0792c dimer. Upper two panels highlight how the C-terminal tail of one chain latches on to the other to stabilize dimeric association. Lower four panels show calculated collective motion in the protein structure which shows motion in the N-termini of the protein chains in the dimer.

**Figure S7:**
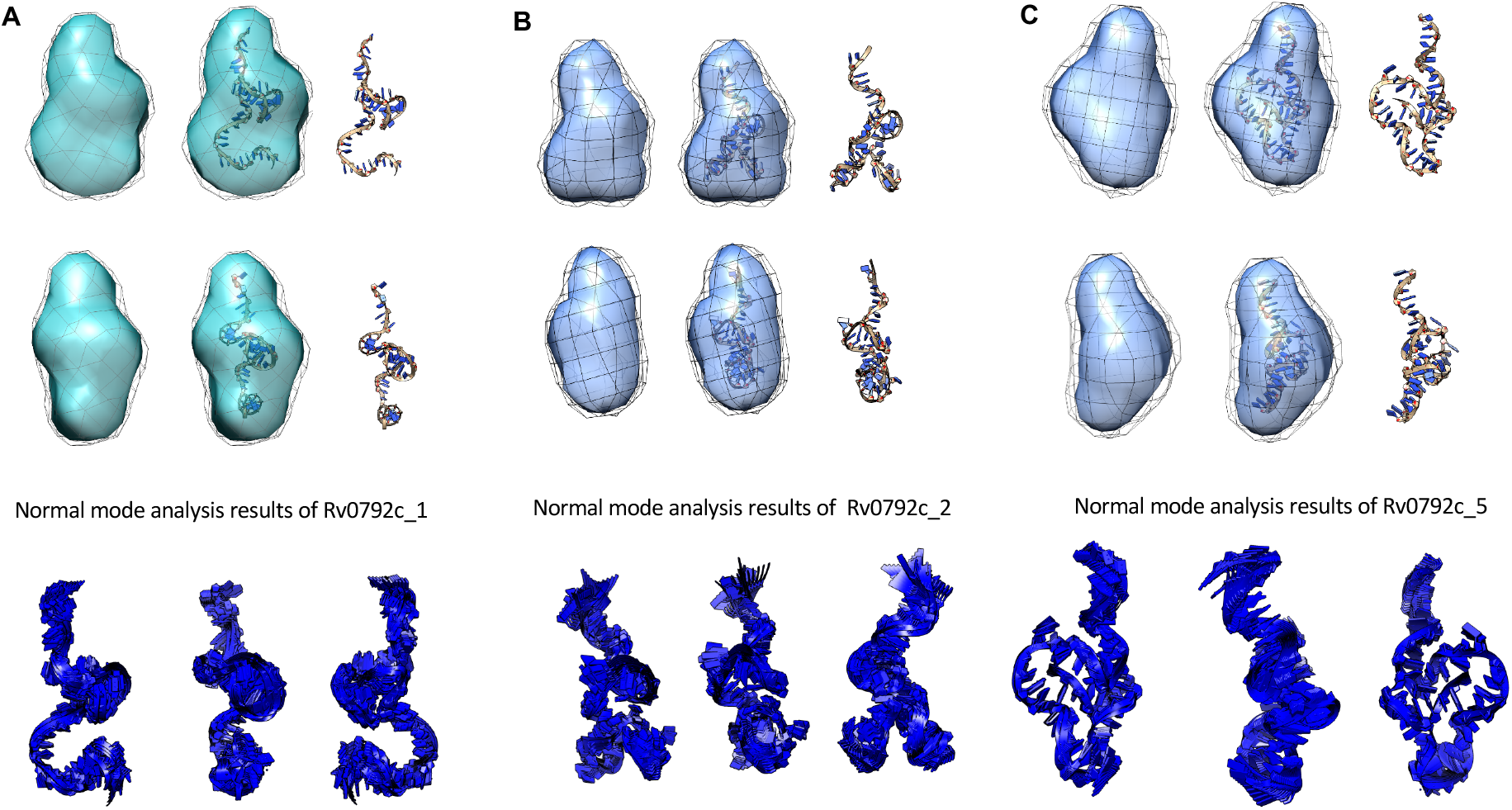
The SAXS based shapes of the three ssDNA aptamers have been shown here. The maps indicate the shape common to ten independent dummy residue solutions and black mesh represents the variation in them. The right panels show the low energy conformation of aptamer which best resembled the shape profile obtained experimentally in their SAXS profiles. All aptamers are monomer in solution. Normal mode analysis of the residue detail models of the aptamers have been done to perceive the extent of inherent motion accessible to the model of aptamers which also explains to some extent the additional volume seen in the molecular map of scattering shape of aptamers.

**Figure S8:**
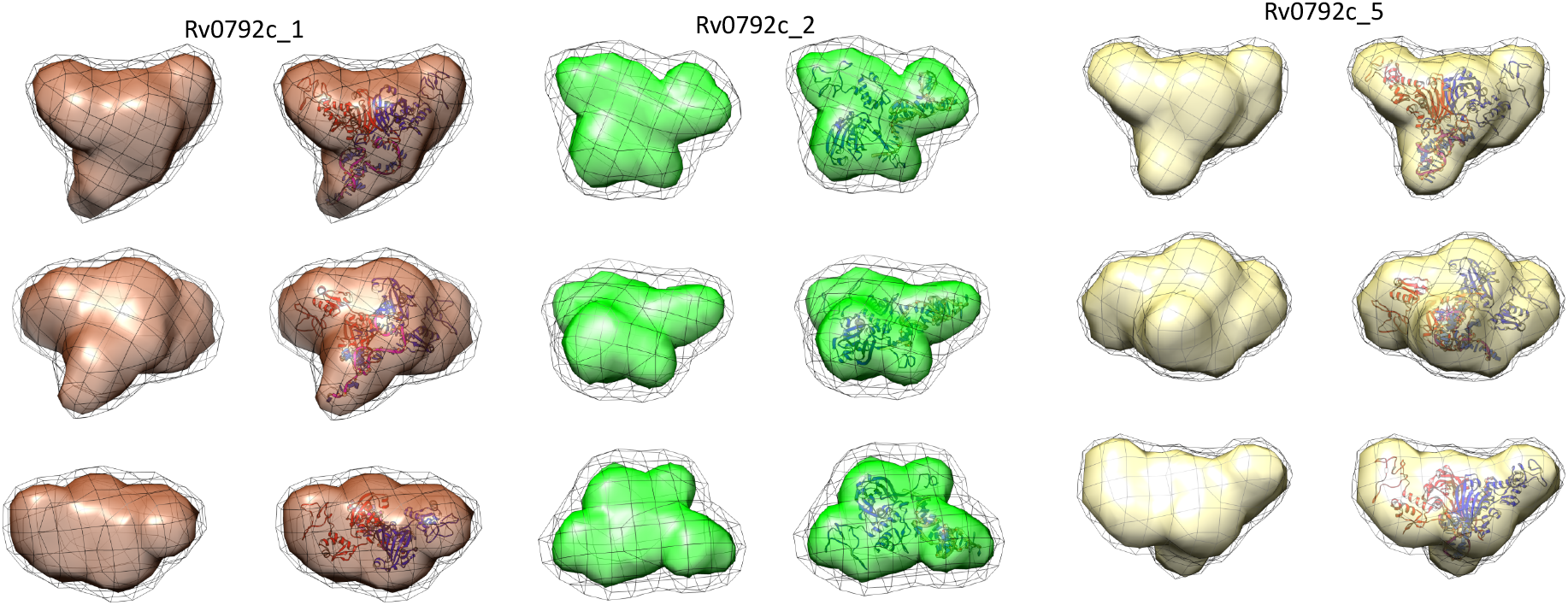
The SAXS based envelope shapes and residue scale models of the three ssDNA aptamers bound with Rv0792c protein have been shown here. The maps indicate the shape common to ten independent dummy residue solutions and black mesh represents the variation in them. The left column of images for the three sets shows the envelope of the complexes. The right column has the residue detailed model of the protein and aptamer where the latter was docked on protein and its pose was selected based on agreement with experimental SAXS data profile. The residue detail model was superimposed on the SAXS data-based envelope in automated manner by aligning their inertial axes.

**Table S1:**
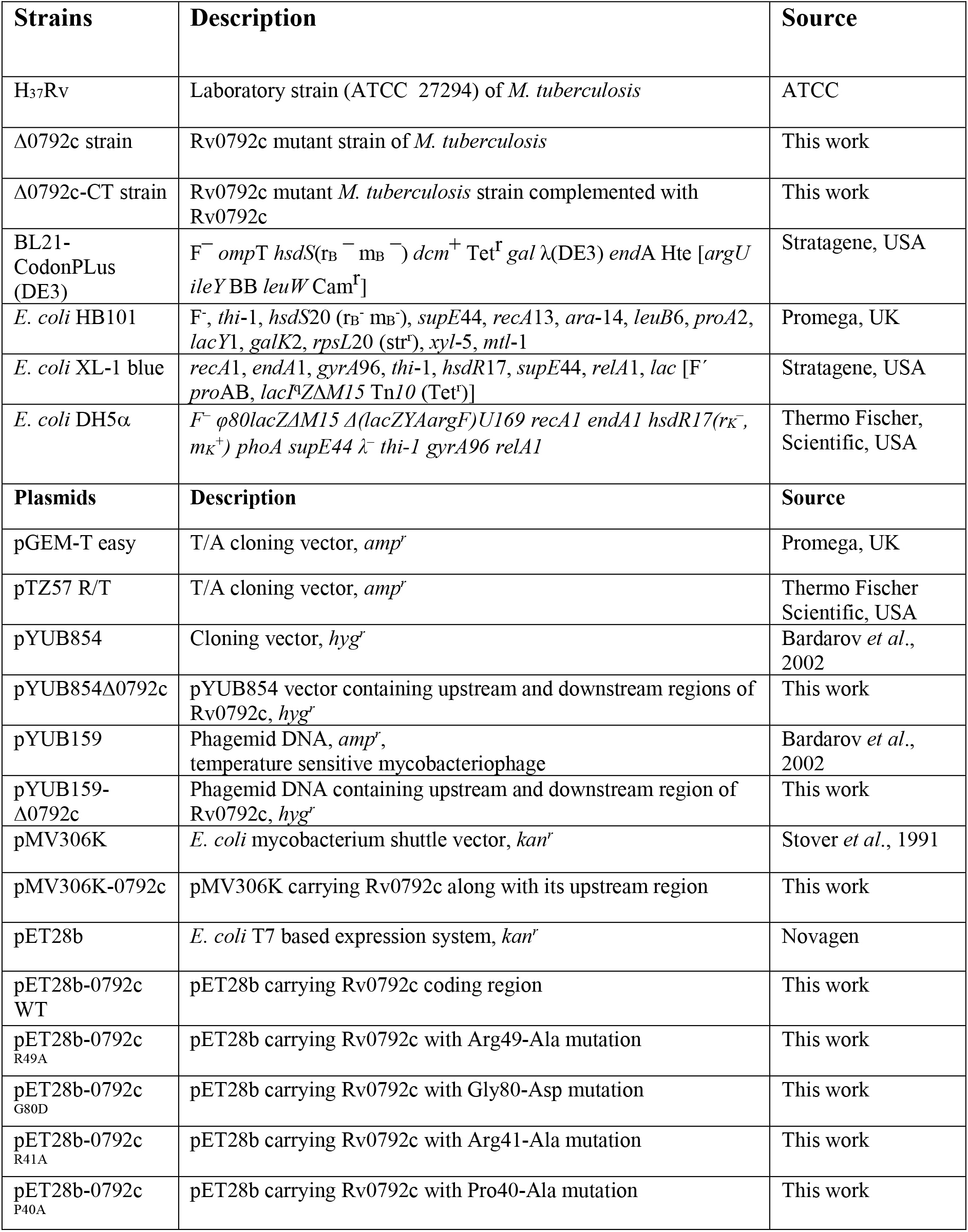
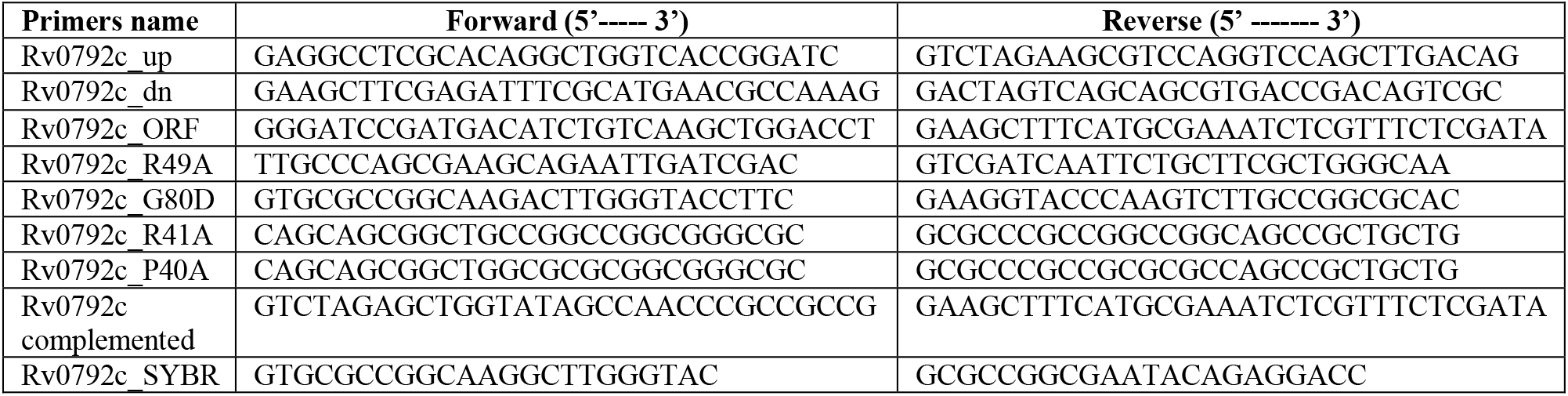
List of strains, plasmids and primers used in the study.

**Table S2:**
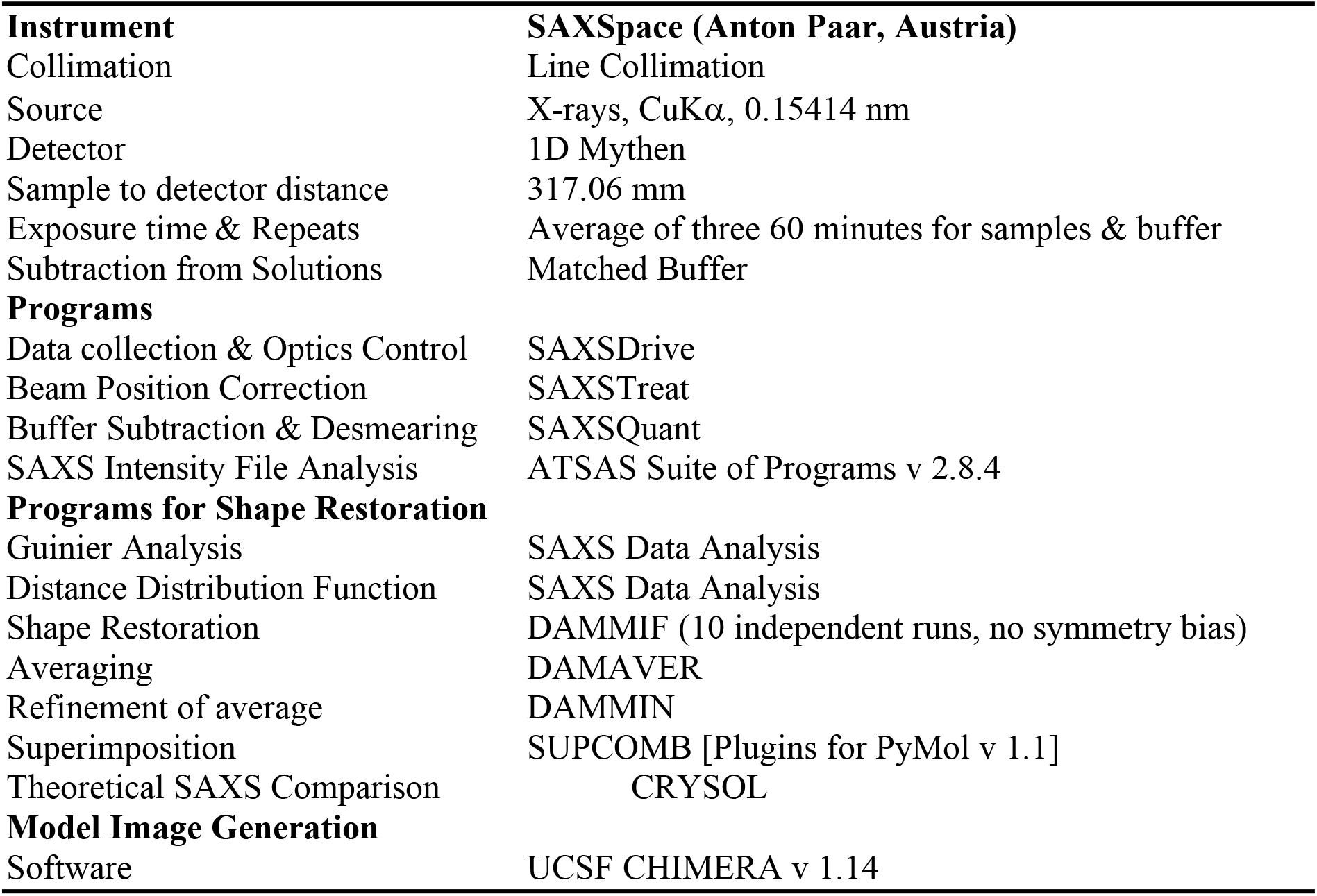
Details of instrumentation, programs used for SAXS processing and SAXS data based parameters of unliganded protein Rv0792c, aptamers found to bind the protein and their molar mixtures are tabulated below.

**Table S3.** SAXS Data Analysis and Shape Reconstruction Related Parameters.

| 1200 | Sample | Guinier | Distance Distribution | | NSD (Models) | $\chi^2$ to |
| --- | --- | --- | --- | --- | --- | --- |
| 1201 |  | Analysis | Function |  |  | residue model |
| 1202 |  | R<sub>g</sub> (nm) | D<sub>max</sub> (nm) | R<sub>g</sub>(nm) |  |  |
| 1203 |  |  |  |  |  |  |
| 1204 | <u>Unliganded Protein</u> |  |  |  |  |  |
| 1205 | GntR | 3.30 | 12.5 | 3.31 | 0.93±0.2 (10) | 1.3 |
| 1206 |  |  |  |  |  |  |
| 1207 | <u>Unliganded Aptamers</u> |  |  |  |  |  |
| 1208 | Rv0792c_5 | 1.81 | 7.1 | 1.84 | 0.549±0.02 (10) | 1.2 |
| 1209 | Rv0792c_1 | 2.10 | 8.6 | 2.23 | 0.727±0.03 (10) | 1.7 |
| 1210 | Rv0792c_2 | 1.92 | 7.6 | 2.01 | 0.684±0.02 (10) | 1.5 |
| 1211 |  |  |  |  |  |  |
| 1212 | <u>1:1 molar mixtures</u> |  |  |  |  |  |
| 1213 | P+ Rv0792c_5 | 3.33 | 12.5 | 3.40 | 0.92±0.1 (10) | 1.7 |
| 1214 | P+ Rv0792c_1 | 3.52 | 12.4 | 3.55 | 0.84±0.2 (10) | 1.8 |
| 1215 | P+ Rv0792c_2 | 3.10 | 9.8 | 3.08 | 0.78±0.1 (10) | 1 |

## Acknowledgements

RS acknowledges the funding received from Department of Biotechnology, India (Grant ID; BT/PR30215/MED/29/1343/2018). TKS thanks Department of Biotechnology, India for funding support through Translational Research Program (BT/PR30159/MED/15/188/2018). Financial Support provided to NKC from the DST SERB-NPDF Program (PDF/2016/002392) is gratefully acknowledged. AS acknowledges research fellowship received from Indian Council of Medical Research. TPG is also thankful to Department of Biotechnology for her fellowship. The authors are also thankful to Infection disease research facility and small animal house staff members at THSTI for technical help. RS is a recipient of Ramalingaswami fellowship and National Bioscience Award from Department of Biotechnology. RS is a senior fellow of Wellcome Trust-DBT India Alliance. The authors acknowledge lab attendants Mr. Rajesh and Mr. Sher Singh for technical help.

## Author contributions

RS and TKS conceived and designed the work plan. NKC, AS and TPG performed cloning, biochemical, microbiology and animal experiments. AA performed SELEX based experiments. KD and A performed SAXS studies. EK and AK performed CD studies. PS performed AUC studies. RS, TKS, A, DS and AK supervised the experiments performed in their respective laboratories. RS, TKS, A and NKC analyzed the data, interpreted them, and wrote the paper as well.

## Conflicts of interest statement

The authors declare that the research was conducted in the absence of any commercial or financial relationships that could be considered as a potential conflict of interest.

## Notes

### Competing Interest Statement

The authors have declared no competing interest.

## References

1. Donoghue HD, Spigelman M, Greenblatt CL, Lev-Maor G, Bar-Gal GK, Matheson C, et al. Tuberculosis: from prehistory to Robert Koch, as revealed by ancient DNA. Lancet Infect Dis. 2004;4(9):584–92.

2. Stewart GR, Robertson BD, Young DB. Tuberculosis: a problem with persistence. Nat Rev Microbiol. 2003;1(2):97–105.

3. Zumla A, Abubakar I, Raviglione M, Hoelscher M, Ditiu L, McHugh TD, et al. Drug-resistant tuberculosis--current dilemmas, unanswered questions, challenges, and priority needs. J Infect Dis. 2012;205 Suppl 2:S228–40.

4. Kliiman K, Altraja A. Predictors of poor treatment outcome in multi- and extensively drug-resistant pulmonary TB. Eur Respir J. 2009;33(5):1085–94.

5. Farley JE, Ram M, Pan W, Waldman S, Cassell GH, Chaisson RE, et al. Outcomes of multi-drug resistant tuberculosis (MDR-TB) among a cohort of South African patients with high HIV prevalence. PLoS One. 2011;6(7):e20436.

6. Brandt L, Feino Cunha J, Weinreich Olsen A, Chilima B, Hirsch P, Appelberg R, et al. Failure of the Mycobacterium bovis BCG vaccine: some species of environmental mycobacteria block multiplication of BCG and induction of protective immunity to tuberculosis. Infect Immun. 2002;70(2):672–8.

7. Gupta S, Chatterji D. Stress responses in mycobacteria. IUBMB Life. 2005;57(3):149–59.

8. Zahrt TC, Deretic V. Mycobacterium tuberculosis signal transduction system required for persistent infections. Proc Natl Acad Sci U S A. 2001;98(22):12706–11.

9. Flentie K, Garner AL, Stallings CL. Mycobacterium tuberculosis Transcription Machinery: Ready To Respond to Host Attacks. J Bacteriol. 2016;198(9):1360–73.

10. Rigali S, Derouaux A, Giannotta F, Dusart J. Subdivision of the helix-turn-helix GntR family of bacterial regulators in the FadR, HutC, MocR, and YtrA subfamilies. J Biol Chem. 2002;277(15):12507–15.

11. Perez-Rueda E, Hernandez-Guerrero R, Martinez-Nunez MA, Armenta-Medina D, Sanchez I, Ibarra JA. Abundance, diversity and domain architecture variability in prokaryotic DNA-binding transcription factors. PLoS One. 2018;13(4):e0195332.

12. Haydon DJ, Guest JR. A new family of bacterial regulatory proteins. FEMS Microbiol Lett. 1991;63(2-3):291–5.

13. Suvorova IA, Korostelev YD, Gelfand MS. GntR Family of Bacterial Transcription Factors and Their DNA Binding Motifs: Structure, Positioning and Co-Evolution. PLoS One. 2015;10(7):e0132618.

14. van Aalten DM, DiRusso CC, Knudsen J. The structural basis of acyl coenzyme A-dependent regulation of the transcription factor FadR. EMBO J. 2001;20(8):2041–50.

15. Jaques S, McCarter LL. Three new regulators of swarming in Vibrio parahaemolyticus. J Bacteriol. 2006;188(7):2625–35.

16. Hoskisson PA, Rigali S. Chapter 1: Variation in form and function the helix-turn-helix regulators of the GntR superfamily. Adv Appl Microbiol. 2009;69:1–22.

17. Hillerich B, Westpheling J. A new GntR family transcriptional regulator in streptomyces coelicolor is required for morphogenesis and antibiotic production and controls transcription of an ABC transporter in response to carbon source. J Bacteriol. 2006;188(21):7477–87.

18. Haine V, Sinon A, Van Steen F, Rousseau S, Dozot M, Lestrate P, et al. Systematic targeted mutagenesis of Brucella melitensis 16M reveals a major role for GntR regulators in the control of virulence. Infect Immun. 2005;73(9):5578–86.

19. Biswas RK, Dutta D, Tripathi A, Feng Y, Banerjee M, Singh BN. Identification and characterization of Rv0494: a fatty acid-responsive protein of the GntR/FadR family from Mycobacterium tuberculosis. Microbiology. 2013;159(Pt 5):913–23.

20. Poupin P, Ducrocq V, Hallier-Soulier S, Truffaut N. Cloning and characterization of the genes encoding a cytochrome P450 (PipA) involved in piperidine and pyrrolidine utilization and its regulatory protein (PipR) in Mycobacterium smegmatis mc2155. J Bacteriol. 1999;181(11):3419–26.

21. Casali N, White AM, Riley LW. Regulation of the Mycobacterium tuberculosis mce1 operon. J Bacteriol. 2006;188(2):441–9.

22. Santangelo Mde L, Blanco F, Campos E, Soria M, Bianco MV, Klepp L, et al. Mce2R from Mycobacterium tuberculosis represses the expression of the mce2 operon. Tuberculosis (Edinb). 2009;89(1):22–8.

23. Zeng J, Deng W, Yang W, Luo H, Duan X, Xie L, et al. Mycobacterium tuberculosis Rv1152 is a Novel GntR Family Transcriptional Regulator Involved in Intrinsic Vancomycin Resistance and is a Potential Vancomycin Adjuvant Target. Sci Rep. 2016;6:28002.

24. Singh R, Singh M, Arora G, Kumar S, Tiwari P, Kidwai S. Polyphosphate deficiency in Mycobacterium tuberculosis is associated with enhanced drug susceptibility and impaired growth in guinea pigs. J Bacteriol. 2013;195(12):2839–51.

25. Kidwai S, Park CY, Mawatwal S, Tiwari P, Jung MG, Gosain TP, et al. Dual Mechanism of Action of 5-Nitro-1,10-Phenanthroline against Mycobacterium tuberculosis. Antimicrob Agents Chemother. 2017;61(11).

26. Bardarov S, Bardarov S, Jr., Pavelka MS, Jr., Sambandamurthy V, Larsen M, Tufariello J, et al. Specialized transduction: an efficient method for generating marked and unmarked targeted gene disruptions in Mycobacterium tuberculosis, M. bovis BCG and M. smegmatis. Microbiology. 2002;148(Pt 10):3007–17.

27. Agarwal S, Sharma A, Bouzeyen R, Deep A, Sharma H, Mangalaparthi KK, et al. VapBC22 toxin-antitoxin system from Mycobacterium tuberculosis is required for pathogenesis and modulation of host immune response. Sci Adv. 2020;6(23):eaba6944.

28. Deep A, Tiwari P, Agarwal S, Kaundal S, Kidwai S, Singh R, et al. Structural, functional and biological insights into the role of Mycobacterium tuberculosis VapBC11 toxin-antitoxin system: targeting a tRNase to tackle mycobacterial adaptation. Nucleic Acids Res. 2018;46(21):11639–55.

29. Kalra P, Mishra SK, Kaur S, Kumar A, Prasad HK, Sharma TK, et al. G-Quadruplex-Forming DNA Aptamers Inhibit the DNA-Binding Function of HupB and Mycobacterium tuberculosis Entry into Host Cells. Mol Ther Nucleic Acids. 2018;13:99–109.

30. Anil Kumar V, Goyal R, Bansal R, Singh N, Sevalkar RR, Kumar A, et al. EspR-dependent ESAT-6 Protein Secretion of Mycobacterium tuberculosis Requires the Presence of Virulence Regulator PhoP. J Biol Chem. 2016;291(36):19018–30.

31. Waterhouse A, Bertoni M, Bienert S, Studer G, Tauriello G, Gumienny R, et al. SWISS-MODEL: homology modelling of protein structures and complexes. Nucleic Acids Res. 2018;46(W1):W296–W303.

32. Pandey K, Rathore YS, Nath SK, Ashish. Towards strain-independent anti-influenza peptides: a SAXS- and modeling-based study. Journal of biomolecular structure & dynamics. 2013.

33. Gautam JK, Ashish, Comeau LD, Krueger JK, Smith MF. Structural and functional evidence for the role of the TLR2 DD loop in TLR1/TLR2 heterodimerization and signaling. J Biol Chem. 2006;281(40):30132–42.

34. Rathore YS, Rehan M, Pandey K, Sahni G, Ashish. First structural model of full-length human tissue-plasminogen activator: a SAXS data-based modeling study. The journal of physical chemistry B. 2012;116(1):496–502.

35. Suhre K, Navaza J, Sanejouand YH. NORMA: a tool for flexible fitting of high- resolution protein structures into low-resolution electron-microscopy-derived density maps. Acta crystallographica Section D, Biological crystallography. 2006;62(Pt 9):1098–100.

36. Sampaio MM, Chevance F, Dippel R, Eppler T, Schlegel A, Boos W, et al. Phosphotransferase-mediated transport of the osmolyte 2-O-alpha-mannosyl-D-glycerate in Escherichia coli occurs by the product of the mngA (hrsA) gene and is regulated by the mngR (farR) gene product acting as repressor. J Biol Chem. 2004;279(7):5537–48.

37. Fillenberg SB, Grau FC, Seidel G, Muller YA. Structural insight into operator dre-sites recognition and effector binding in the GntR/HutC transcription regulator NagR. Nucleic Acids Res. 2015;43(2):1283–96.

38. Fillenberg SB, Friess MD, Korner S, Bockmann RA, Muller YA. Crystal Structures of the Global Regulator DasR from Streptomyces coelicolor: Implications for the Allosteric Regulation of GntR/HutC Repressors. PLoS One. 2016;11(6):e0157691.

39. Vasanthakumar A, Kattusamy K, Prasad R. Regulation of daunorubicin biosynthesis in Streptomyces peucetius - feed forward and feedback transcriptional control. J Basic Microbiol. 2013;53(8):636–44.

40. Martinez-Antonio A, Collado-Vides J. Identifying global regulators in transcriptional regulatory networks in bacteria. Curr Opin Microbiol. 2003;6(5):482–9.

41. Raman N, Black PN, DiRusso CC. Characterization of the fatty acid-responsive transcription factor FadR. Biochemical and genetic analyses of the native conformation and functional domains. J Biol Chem. 1997;272(49):30645–50.

42. Ehrt S, Schnappinger D. Mycobacterial survival strategies in the phagosome: defence against host stresses. Cell Microbiol. 2009;11(8):1170–8.

43. Pinheiro J, Lisboa J, Pombinho R, Carvalho F, Carreaux A, Brito C, et al. MouR controls the expression of the Listeria monocytogenes Agr system and mediates virulence. Nucleic Acids Res. 2018;46(18):9338–52.

44. Gu D, Meng H, Li Y, Ge H, Jiao X. A GntR Family Transcription Factor (VPA1701) for Swarming Motility and Colonization of Vibrio parahaemolyticus. Pathogens. 2019;8(4).

45. Li Z, Xiang Z, Zeng J, Li Y, Li J. A GntR Family Transcription Factor in Streptococcus mutans Regulates Biofilm Formation and Expression of Multiple Sugar Transporter Genes. Front Microbiol. 2018;9:3224.

46. Chai Y, Kolter R, Losick R. A widely conserved gene cluster required for lactate utilization in Bacillus subtilis and its involvement in biofilm formation. J Bacteriol. 2009;191(8):2423–30.

47. Park HD, Guinn KM, Harrell MI, Liao R, Voskuil MI, Tompa M, et al. Rv3133c/dosR is a transcription factor that mediates the hypoxic response of Mycobacterium tuberculosis. Mol Microbiol. 2003;48(3):833–43.

48. Betts JC, Lukey PT, Robb LC, McAdam RA, Duncan K. Evaluation of a nutrient starvation model of Mycobacterium tuberculosis persistence by gene and protein expression profiling. Mol Microbiol. 2002;43(3):717–31.

49. Singh A, Jain S, Gupta S, Das T, Tyagi AK. mymA operon of Mycobacterium tuberculosis: its regulation and importance in the cell envelope. FEMS Microbiol Lett. 2003;227(1):53–63.

50. Singh A, Gupta R, Vishwakarma RA, Narayanan PR, Paramasivan CN, Ramanathan VD, et al. Requirement of the mymA operon for appropriate cell wall ultrastructure and persistence of Mycobacterium tuberculosis in the spleens of guinea pigs. J Bacteriol. 2005;187(12):4173–86.

51. Dhiman A, Haldar S, Mishra SK, Sharma N, Bansal A, Ahmad Y, et al. Generation and application of DNA aptamers against HspX for accurate diagnosis of tuberculous meningitis. Tuberculosis (Edinb). 2018;112:27–36.

52. Schroder J, Maus I, Ostermann AL, Kogler AC, Tauch A. Binding of the IclR-type regulator HutR in the histidine utilization (hut) gene cluster of the human pathogen Corynebacterium resistens DSM 45100. FEMS Microbiol Lett. 2012;331(2):136–43.

53. Bender RA. Regulation of the histidine utilization (hut) system in bacteria. Microbiol Mol Biol Rev. 2012;76(3):565–84.

54. Aravind L, Anantharaman V. HutC/FarR-like bacterial transcription factors of the GntR family contain a small molecule-binding domain of the chorismate lyase fold. FEMS Microbiol Lett. 2003;222(1):17–23.

55. Sieira R, Arocena GM, Bukata L, Comerci DJ, Ugalde RA. Metabolic control of virulence genes in Brucella abortus: HutC coordinates virB expression and the histidine utilization pathway by direct binding to both promoters. J Bacteriol. 2010;192(1):217–24.

56. Sharma TK, Bruno JG, Dhiman A. ABCs of DNA aptamer and related assay development. Biotechnol Adv. 2017;35(2):275–301.

57. Chatterjee B, Kalyani N, Anand A, Khan E, Das S, Bansal V, et al. GOLD SELEX: a novel SELEX approach for the development of high-affinity aptamers against small molecules without residual activity. Mikrochim Acta. 2020;187(11):618.

58. Badmalia MD, Sharma P, Yadav SPS, Singh S, Khatri N, Garg R, et al. Bonsai Gelsolin Survives Heat Induced Denaturation by Forming beta-Amyloids which Leach Out Functional Monomer. Scientific reports. 2018;8(1):12602.

59. Jain D. Allosteric control of transcription in GntR family of transcription regulators: A structural overview. IUBMB Life. 2015;67(7):556–63.

60. DiRusso CC, Black PN, Weimar JD. Molecular inroads into the regulation and metabolism of fatty acids, lessons from bacteria. Prog Lipid Res. 1999;38(2):129–97.

61. Franco IS, Mota LJ, Soares CM, de Sa-Nogueira I. Functional domains of the Bacillus subtilis transcription factor AraR and identification of amino acids important for nucleoprotein complex assembly and effector binding. J Bacteriol. 2006;188(8):3024–36.

62. Resch M, Schiltz E, Titgemeyer F, Muller YA. Insight into the induction mechanism of the GntR/HutC bacterial transcription regulator YvoA. Nucleic Acids Res. 2010;38(7):2485–97.

63. Li ZQ, Zhang JL, Xi L, Yang GL, Wang SL, Zhang XG, et al. Deletion of the transcriptional regulator GntR down regulated the expression of Genes Related to Virulence and Conferred Protection against Wild-Type Brucella Challenge in BALB/c Mice. Mol Immunol. 2017;92:99–105.

64. Zhou X, Yan Q, Wang N. Deciphering the regulon of a GntR family regulator via transcriptome and ChIP-exo analyses and its contribution to virulence in Xanthomonas citri. Mol Plant Pathol. 2017;18(2):249–62.

65. Wang T, Qi Y, Wang Z, Zhao J, Ji L, Li J, et al. Coordinated regulation of anthranilate metabolism and bacterial virulence by the GntR family regulator MpaR in Pseudomonas aeruginosa. Mol Microbiol. 2020;114(5):857–69.

66. Zhou Y, Nie R, Liu X, Kong J, Wang X, Li J. GntR is involved in the expression of virulence in strain Streptococcus suis P1/7. FEMS Microbiol Lett. 2018;365(14).

67. Lemieux MJ, Ference C, Cherney MM, Wang M, Garen C, James MN. The crystal structure of Rv0793, a hypothetical monooxygenase from M. tuberculosis. J Struct Funct Genomics. 2005;6(4):245–57.

68. Jing M, Bowser MT. Methods for measuring aptamer-protein equilibria: a review. Anal Chim Acta. 2011;686(1-2):9–18.

69. Bilibana MP, Citartan M, Yeoh TS, Rozhdestvensky TS, Tang TH. Aptamers as the Agent in Decontamination Assays (Apta-Decontamination Assays): From the Environment to the Potential Application In Vivo. J Nucleic Acids. 2017;2017:3712070.

70. Taneja V, Goel M, Shankar U, Kumar A, Khilnani GC, Prasad HK, et al. An Aptamer Linked Immobilized Sorbent Assay (ALISA) to Detect Circulatory IFN-alpha, an Inflammatory Protein among Tuberculosis Patients. ACS Comb Sci. 2020;22(11):656–66.

71. Manochehry S, McConnell EM, Li Y. Unraveling Determinants of Affinity Enhancement in Dimeric Aptamers for a Dimeric Protein. Sci Rep. 2019;9(1):17824.

72. Hu J, Kim J, Easley CJ. Quantifying Aptamer-Protein Binding via Thermofluorimetric Analysis. Anal Methods. 2015;7(17):7358–62.

73. Momose I, Kunimoto S, Osono M, Ikeda D. Inhibitors of insulin-like growth factor-1 receptor tyrosine kinase are preferentially cytotoxic to nutrient-deprived pancreatic cancer cells. Biochem Biophys Res Commun. 2009;380(1):171–6.

74. Davis MI, Sasaki AT, Shen M, Emerling BM, Thorne N, Michael S, et al. A homogeneous, high-throughput assay for phosphatidylinositol 5-phosphate 4-kinase with a novel, rapid substrate preparation. PLoS One. 2013;8(1):e54127.

